# Axons organize into micro-tracts around the cortical vasculature

**DOI:** 10.64898/2026.09.26.754479

**Authors:** Javier J How, Bethanny Danskin, Benjamin D Pedigo, Elana M Memke, Stefan Stamenkovic, David Kleinfeld, Andy Y Shih

## Abstract

Neural activity is metabolically costly and supported by local increases in cerebral blood flow. The process by which neural activity drives blood flow increase, neurovascular coupling (NVC), remains poorly understood. One persistent knowledge gap is how subcellular compartments are arranged at the vascular wall, where local signaling occurs. This requires a survey of the neurovascular interface at the nanometer scale. To this end, we analyzed a ∼1 mm^3^ volume of mouse visual cortex imaged with serial electron microscopy. We manually labeled different vascular zones and classified the EM segments surrounding hundreds of vessel segments across these zones. We found that capillaries and venous vessels are surrounded by more axonal volume than arteriolar vessels. Perivascular axons tended to bundle into micro-tracts, where axons coursed near the vessel wall in a variety of geometric orientations. Micro-tracts covered ∼60% of the surface of capillaries, but only 24 – 40% of every other vascular zone’s surface. Finally, we show that the basket cells, with their extensive axonal arbors, are candidate members of micro-tracts: on average, their axons approach 2-to-3-fold more capillaries than other neuronal subtypes. Altogether, perivascular axonal micro-tracts may be an important physical substrate for NVC signaling at capillaries.

## Introduction

Increases in neuronal activity induce a local upregulation of cerebral blood flow (CBF) through a process referred to as neurovascular coupling (NVC). This coupling is thought to supply glucose and oxygen to active tissue, clear metabolic waste, and regulate brain temperature (Iadecola, 2017).

During NVC, neurons and astrocytes release potassium ions, nitric oxide, arachidonic acid derivatives, and other vasoactive agents onto the endothelial cells, pericytes, and smooth muscle cells (SMCs) that form the vascular wall (Iadecola, 2017). These signals act upon specific zones with the vascular network, necessitating a deeper understanding of how the neurovascular interface differs along the vasculature. In the cerebral cortex, pial arterioles at the brain surface branch and descend as penetrating arterioles (PA) and pass through different cortical layers. SMCs on the PA hyperpolarize and relax, leading to vasodilation and blood flow increases throughout the downstream capillary network (Filosa et al., 2006). Ensheathing pericytes on the arteriole-capillary transition zone are also dynamic participants in blood flow regulation, while capillary pericytes further downstream modulate capillary diameter on a slower timescale (Grant et al., 2019; Hall et al., 2014; Hamilton et al., 2010; Hartmann et al., 2022).

There are at least two different vessel-type-specific mechanisms by which neurons can trigger increases in CBF. One model emphasizes capillaries as loci of NVC. Capillaries account for most of the length of the cortical vascular network and a typical neuron lies ∼15 μm from the nearest one (Ji et al., 2021; Tsai et al., 2009), so they are well-positioned to sample neural activity at a much finer spatial scale than the sparsely spaced PAs. Longden et al. (2017) (Figure 1A) showed that capillary endothelial cells are hyperpolarized by modest rises in extracellular K⁺, and that this hyperpolarization retrogradely propagates along the endothelium to the upstream PA, where it hyperpolarizes SMCs and dilates the PA feeding the active region. Consistent with this, the dilation of capillaries near active neurons can precede dilation of the upstream arteriole (Hall et al., 2014). In this model, the capillary is a primary sensor of neuronal activity, and it is sensitive to K⁺ released during the propagation of action potentials and synaptic transmission. That implies a specific anatomy: the neuronal structures that release K⁺ – axons and synapses – should lie near capillaries, close enough for the release of K⁺ to be detected by the capillary endothelium. In a second model, Zhang et al. (2024) found that glutamatergic axons in mouse barrel cortex form direct synapses, through holes in astrocytic endfeet, onto SMCs on PAs. This suggests a second anatomy: axons form synapses, or at least directly touch, the mural cells on PAs.

**Figure 1.**
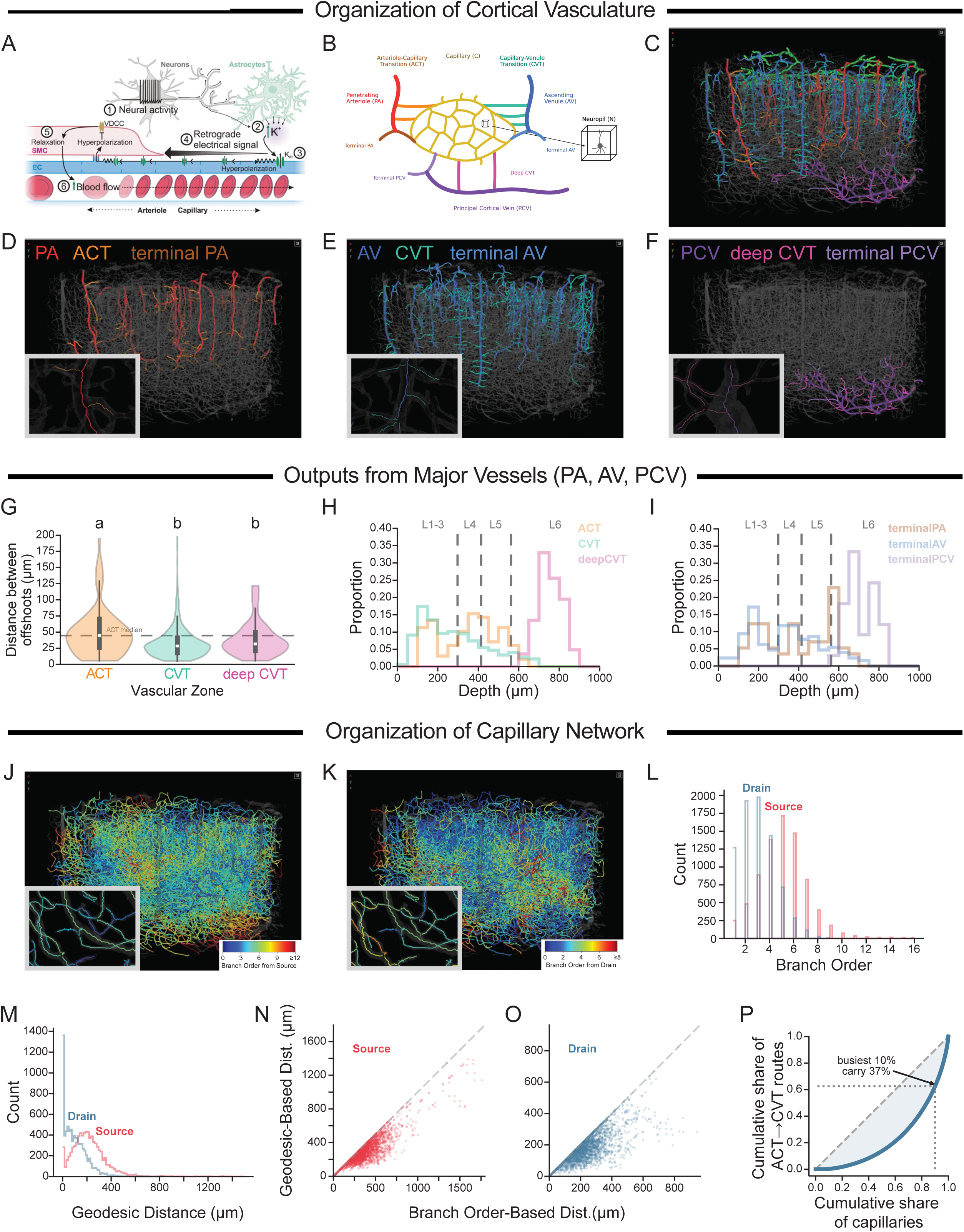
The organization of the vasculature in the MICrONs mouse visual cortex. (A) Model of neurovascular coupling from Longden et al. (2017). (B) Schematic of canonical cortical vasculature. (C) Overview of all non-capillary blood vessels in the MICrONS dataset. (D) Blood sources in the brain: penetrating arterioles (PA, n = 21; red) supply blood that enter the capillary network via the arteriole-capillary transition zones (ACTs, n = 99; bright orange) and terminal PAs (dark orange; n = 57 red). Inset shows one PA (red) and two ACTs (bright orange). (D, E) Blood drains from the brain via two systems: (D) capillary-venule transition zones (CVTs, n = 593; turquoise) and terminal ascending venules (terminal AVs, n = 128; steel blue) through AVs (n = 46; royal blue), as well as (E) deep CVTs (n = 82, magenta-pink) and terminal principal cortical veins (terminal PCVs, n = 33; lavender) through PCVs (n = 2; deep violet). Insets depict (E) one AV (royal blue) and 5 CVTs (turquoise) and (F) one PCV (deep violet) and four deep CVTs (magenta-pink). (G) Distribution of distances between ACTs, CVTs, and deep CVTs. Y-axis was limited to 200 μm; thus, 8% of ACTs and 0.6% of CVTs are not shown. Violins show the distribution, the box the interquartile range and the white line the median. Vascular zones sharing a letter are not significantly different; those sharing no letter differ at adjusted p < 0.05 (Kruskal–Wallis p = 4.72e-07; Dunn’s post-hoc with Bonferonni correction: ACT vs CVT p = 5.90e-07, and ACT vs deep CVT p = 0.035). (H, I) Distribution of (H) transition zones and (I) terminal zones as a function of cortical depth. (J, K) Capillaries (n = 7757 out of a total of 9292 capillaries) colored according to their branch order from (J) blood source or (K) blood drain; blue means closer, red means farther. Inset shows several capillaries at varying number of branches from the nearest source or drain. (L) Distribution of capillary branch order from nearest blood source (red; ACT or terminal PA) or drain (blue; CVT, deep CVT, terminal AV, or terminal PCV). (M) Like L, but using geodesic distance. (N, O) Branch order vs geodesic distance for (N) sources and (O) drains. (P) Traffic through the capillary bed is unevenly distributed. Capillaries ranked by the number of ACT→CVT shortest paths passing through them (n = 7,757; 3.8% do not lie on a shortest path). The busiest 10% carry 37.5% of all path traversals (Gini = 0.54); the dashed diagonal line is uniform traffic. Data in C – F, J, and K can be seen at https://spelunker.cave-explorer.org/#!middleauth+https://global.daf-apis.com/nglstate/api/v1/5249453187923968.

These models make specific predictions: 1) axon organization around capillaries must enable the local concentration of K^+^ to reach a high level, and 2) contacts between axons and blood vessels should occur at the PA, even in other brain regions. To this end, we characterized the ultrastructural anatomy (imaged with electron microscopy (EM) at a resolution of 8 × 8 × 40 nm^3^) around blood vessels in a ∼1 mm³ volume of mouse visual cortex (the MICrONS dataset) (MICrONS Consortium, 2025; Wan et al., 2025). The breadth of this large dataset allows us to characterize an unprecedented number of axon-capillary encounters. We manually labeled all branches of the vascular network from arterial inputs to venous drains and then classified EM segments surrounding 347 branches and within 80 neuropil regions as axon, dendrite, soma, thick/myelinated axon, glia, or perivascular components. We asked: (1) what is the heterogeneity in the organization of these components and of synaptic density at the different vascular zones? and (2) is there structure to the micro-organization of axons and dendrites as they encounter vessels?

## Results

### Organization of cortical vasculature in a ∼1 mm^3^ volume of mouse visual cortex

The cortical vasculature comprises the major vessels that source or drain blood, capillaries that spread blood throughout the parenchyma, and connections between the major vessels and capillaries. We skeletonized the EM vascular segmentation (Wan et al., 2025) of the MICrONS dataset and manually labeled the vascular zones. Vertically-oriented penetrating arterioles (PAs) and ascending venules (AVs) source and drain blood to the cortical parenchyma (Figure 1B, D, E). A rarer venous structure, principal cortical veins (PCVs), drain blood primarily from the deeper cortical layers and callosal white matter through long arbors at the gray white matter interface (Figure 1F) (Stamenkovic et al., 2025). We distinguished specific structures relevant to our examination of the microvascular network and developed terminology to identify them (Figure 1B). As PAs penetrate the cortex, they send blood to the capillary network via offshoots called arteriole-capillary-transition zones (ACTs; Figure 1B – D). At their terminal ends, we refer to the offshoots as terminal PAs (Figure 1B – D). Similarly, AVs receive blood from the capillary network via capillary-venule-transition zones (CVTs) and their terminal ends, called terminal AVs, while PCVs drain the capillaries via deep CVTs and terminal PCVs (Figure 1B, C, E, F). The ACTs, CVTs, and deep CVTs tended to emanate from their parent vessels at an ∼90–110° angle, while the terminal branches formed ∼120–140° angles (Supplementary Figure 1G).

Poor trans-cardial perfusion and tissue fixation in EM imaging can lead to collapsed vessels and distort angioarchitecture. However, vessel shape was well preserved in the MICrONS dataset (Sargent et al., 2023). As expected from *in vivo* studies, PAs, AVs, and PCVs were greater in surface area and diameter (Supplementary Figure 1A, B), and less tortuous than capillaries (Supplementary Figure 1D). Surrounding these vessels are densely packed cellular nuclei and processes, all visible at nanoscale resolution. Since tissue shrinkage is also a concern with EM tissue processing, we examined the average neuronal distance from vessels. The median distance between a blood vessel and a non-vascular parenchymal coordinate was 12.9 μm (Supplementary Figure 1E), and the median distance between a neuronal nuclei and the nearest blood vessel was 11.5 μms (Supplementary Figure 1F). This is smaller than the 15 μm distance previously reported in immunostained mouse vibrissa somatosensory cortex, consistent with some tissue shrinkage (Tsai et al., 2009).

This vascular network in the MICrONS dataset contains 21 PAs, 46 AVs, and 2 PCV branches (Figure 1D–F). Our prior analysis reached a different count because it characterized only half the dataset (Bonney et al., 2022), but assessment of the full dataset provides vessel counts more consistent with Grubb (2023). There are many more CVT branches (n = 593) draining into AVs, than ACT offshoots from PAs (n = 99). Similarly, there are more terminal AVs (n = 128) than terminal PAs (n = 57). The ACT branches are spaced a median 44.6 μm apart along the PAs, while the more numerous CVTs and deep CVTs are spaced every 28.4 μm and 32.9 μm, respectively, though the distributions substantially overlap (Dunn’s post-hoc p < 0.05 for both comparisons against ACT; Figure 1G). The ACT branches are spread throughout the layers of cortex, but are scarcer around layers 2 (L2) and 6 (L6) (Figure 1H). Conversely, CVT branches are most common at L1 and reduce in density until they are absent in L6 (Figure 1H). At these depths, the branches of PCVs, deep CVTs, predominate (Figure 1H). Finally, the terminal branches of the major vessels follow a similar pattern as the offshoots (Figure 1I).

We next characterized the topology of the capillary network, focusing on their distance from the nearest blood source (ACT or terminal PA) and drain (CVT, terminal AV, deep CVT, or terminal PCV). The capillary network consists of 9,292 branches, but we focused on the subset of 7,757 branches that were located between a blood source and drain without having to trace a path through a major vessel (Figure 1J, K). Distance was defined in two ways: 1) the number of branches away from the nearest source or drain, where branch order “1” relative to a source meant that the capillary was directly connected to an ACT or terminal PA branch, and 2) the geodesic distance – that is, distance along the blood vessel skeleton – between the capillary and the nearest source or drain, where distance “0 μm” relative to a source meant that the capillary was directly connected to an ACT or terminal PA branch. Because drains are more numerous, each serves a smaller territory: both the branch order and the geodesic distance to the nearest drain (median order 3; median distance 88 μm; blue bars and curve in Figure 1L, M) were lower than those to the nearest source (median order 5; median distance 201 μm; red bars and curve in Figure 1L, M); this was a within-capillary difference of 2 orders and 111 μm, with 74% of capillaries geodesically farther from their source than their drain (Wilcoxon signed-rank, p < 10^-300^; Figure 1L, M).

Branch order is a common way to express a capillary’s position relative to a source or drain, but it does not necessarily track geodesic distance: capillary segments vary several-fold in length and tortuosity (Supplementary Figure 1C, D). To quantify this discrepancy, for each capillary we measured the geodesic distance to the source and drain nearest in branch order and compared it with the distance to the geodesically nearest source and drain. The former is by construction an upper bound on the latter, and the bound held for every capillary (Figure 1N, O). Relative to the source, the two coincided for 54% of capillaries; for the remaining 46%, the branch-order-nearest source lay a median of 38 μm farther away (Figure 1N). Agreement was more frequent relative to the drain, at 68%, but the penalty where it failed was larger: a median of 53 μm across the remaining 32% (Figure 1O).

The inverse comparison behaved differently. For the source or drain identified as geodesically nearest, we counted the branches along that route; this can in principle far exceed the minimum when a capillary reaches its source through a chain of short segments. In practice it rarely did: the counts agreed exactly for 80% of capillaries relative to their source and 75% relative to their drain, and where they differed the geodesic target lay a median of one additional branch away (Supplementary Figure 2A, B). Branch order therefore identifies nearly the same topological neighborhood as geodesic distance, but not the same physical one: a given branch order spans a broad range of distances, so branch order should not be read as a proxy for how far a capillary is from its source or drain.

Finally, we noted that links in a transport network can differ widely in how much traffic they carry, with a minority often lying on a disproportionate share of routes. We asked if individual capillaries similarly differ. For each effective ACT–CVT pair we found the topological shortest path, counting each branch as one step, and tallied traversals per capillary. We found that traversals were distributed unevenly: the busiest 10% of capillaries carried 37.5% of them (Gini = 0.54; Figure 1P). Weighting paths by branch length gave the same result (busiest 10%, 38%; Gini = 0.56). Thus, some capillary branches were central to blood flow.

### Axons are enriched near capillaries and venous vessels

The value of EM lies in the ability to see the cellular milieu in which the blood vessels are embedded. Previously, Dorkenwald et al. (2023) developed SegCLR to generate low-dimensional embeddings of local views of segmented objects in the EM images which could distinguish between neuronal subcompartments, such as soma, axon, and dendrite, and glia. We trained a logistic regression classifier that uses these embeddings to classify segmented objects into one of six categories: axon, dendrite, neuronal soma, thick/myelinated axon (a category composed of both thicker and myelinated axons), glia, and perivascular, the last of which includes the endothelial and mural cells that cover the lumen (Figure 2A). We classified the segmented objects up to 5 μm away from 347 blood vessel branches and separately, as controls, for 80 regions of interest at least ∼13 μm away from the nearest blood vessel, designated neuropil (Supplementary Figure 1E). The blood vessel branches were taken across all layers of cortex, and included 38 PA segments, 41 ACT branches, 110 Capillary branches, 80 CVT branches, and 78 AV segments. Our classifier ignored segmented objects below a size threshold where the embeddings were not generated (Supplementary Figure 3A), and its 5-fold cross-validation accuracy was ≥84% on all labels (Supplementary Figure 3B).

**Figure 2.**
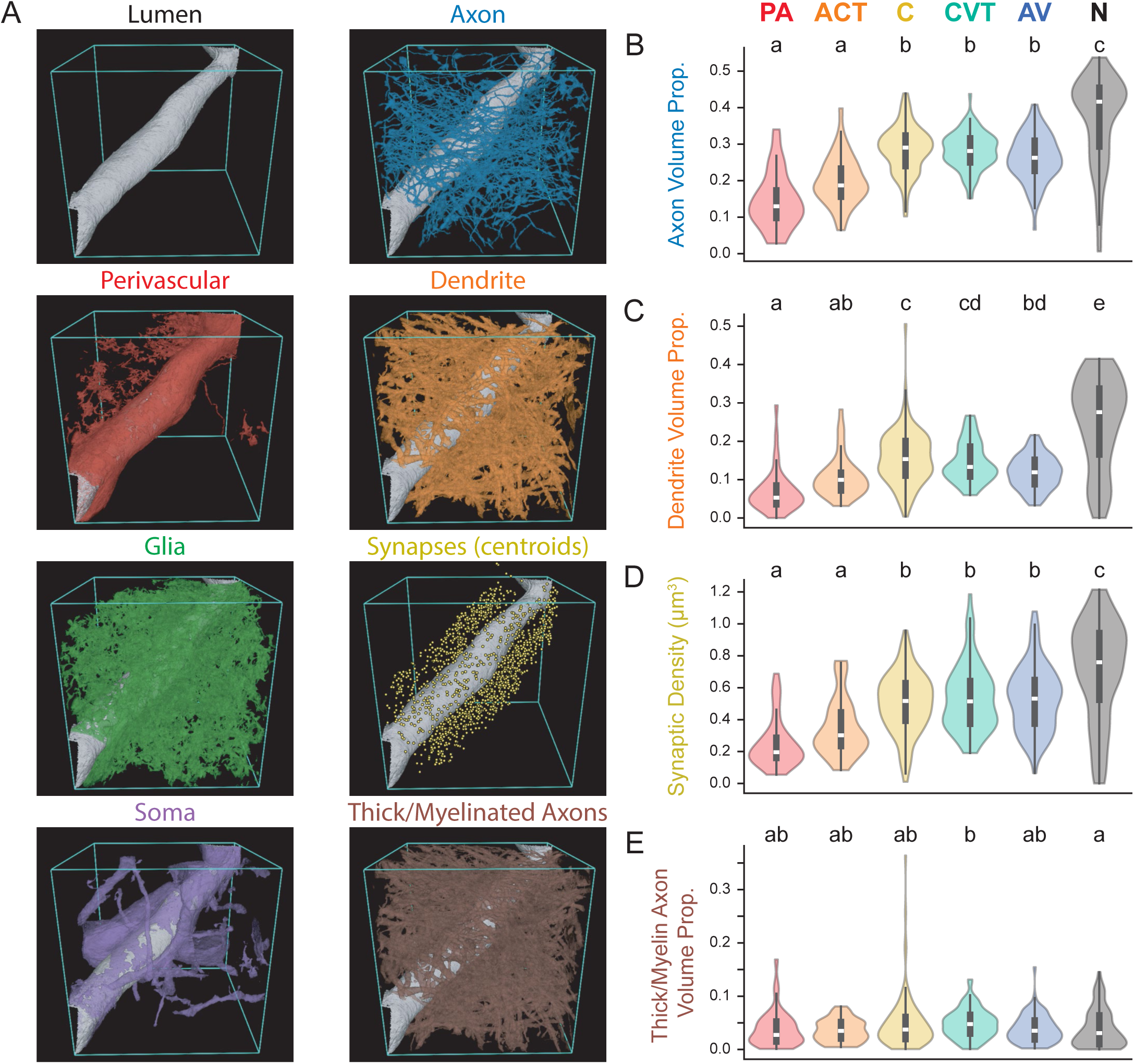
Axons, dendrites, and synapses are enriched near capillaries, CVTs, and AVs. (A) Segments were classified into 1 of 6 categories. Here we show the lumen (grey), all synapses (yellow circles), and only up to 200 segments in each of the six categories; as a result, some synapses are not attached to a depicted axon or dendrite. This example can be seen in https://cj-mesh-bounds-dot-neuroglancer-dot-seung-lab.ue.r.appspot.com/#!middleauth+https://global.daf-apis.com/nglstate/api/v1/5341001120481280. (B, C, E) Proportion of volume within 2 μm of different vascular zones occupied by (B) axons, (C) dendrites, or (E) thick/myelinated axons. (D) Synaptic density within 2 μm of different vascular zones. Violins show the distribution, the box the interquartile range and the white line the median. Vascular zones sharing a letter are not significantly different; those sharing no letter differ at adjusted p < 0.05 (Kruskal–Wallis p = 1.29e-29, 1.95e-24, 1.30e-20, and 0.037 for panels B – E, respectively; Dunn’s post-hoc with Bonferroni correction).

As expected, neuronal compartments were most concentrated in the neuropil (Figure 2B – D). Contrary to a model in which PA and ACT branches are the major sites of neuron-to-vessel communication, we found axons to be more enriched around capillaries, CVT and AV branches relative to both PA and ACT (Dunn’s post-hoc p < 0.05; Figure 2B). Similarly, dendrites occupied more volume around capillaries and CVT branches than near PA and ACT branches (Dunn’s post-hoc p < 0.05; Figure 2C). Synaptic density recapitulated these axon pattern: highest in the neuropil, intermediate around capillaries, CVT and AV, and lowest around PA and ACT (Dunn’s post-hoc p < 0.05; Figure 2D). Thick/myelinated axons and neuronal somata were rare around every vascular zone (median volume proportion 0.03 – 0.05 and 0.02 – 0.05 respectively; Figure 2E, Supplementary Figure 4D). Glia, by contrast, occupied a substantial fraction of every neighborhood and were distributed unevenly: they were least abundant in the neuropil and around PAs, most abundant around capillaries, and intermediate elsewhere (Supplementary Figure 4F). Finally, the perivascular class occupied a larger share of the volume around PAs and ACTs than around capillaries, CVTs or AVs (Supplementary Figure 4G), which may explain the relative lack of neuronal components at those vascular zones.

These results held up at distances up to ∼2 μm from the blood vessels, and by 3 μm away the volume around blood vessels largely resembled the Neuropil (Supplementary Figure 4). Altogether, this suggests that capillaries and the venous vessels are best positioned to integrate local neuronal activity by listening in on signals from axons, dendrites, and synapses.

### Axons bundle into micro-tracts around blood vessels

Grubb (2023) previously surveyed the MICrONS dataset and noted that, in a limited sample of branches, axons wrap around PAs (with perivascular spaces) and nearby capillaries by crossing over them, perpendicular to the direction of the blood vessel. Since we found that axons and dendrites are enriched near capillaries and venous vessels, we sought to determine if axons and dendrites formed these bundles, or micro-tracts, more often near these branches than around PA and ACT branches.

To identify bundles, we clustered axons and dendrites into groups that travel together (Figure 3A, B; Methods). For each vessel branch, the algorithm takes the axons or dendrites within 2 μm of the vessel, trims each to its longest contiguous run inside that shell, computes a direction-aware distance that penalizes pairs whose local directions differ, and clusters them using density-based clustering (Campello et al., 2013); in the neuropil we clustered all axons or dendrites in the analyzed box. In all, we found 6,745 axonal micro-tracts. Axons formed a median of 12 – 21 micro-tracts per branch on every vascular zone (Figure 3C). Strikingly, although axons are most abundant in the neuropil (Figure 2B), they formed almost no micro-tracts there (median 2; Figure 3C), implying that their formation requires a scaffold. Dendrites, on the other hand, rarely formed micro-tracts (median 0–2 across vascular zones; Figure 3D).

**Figure 3.**
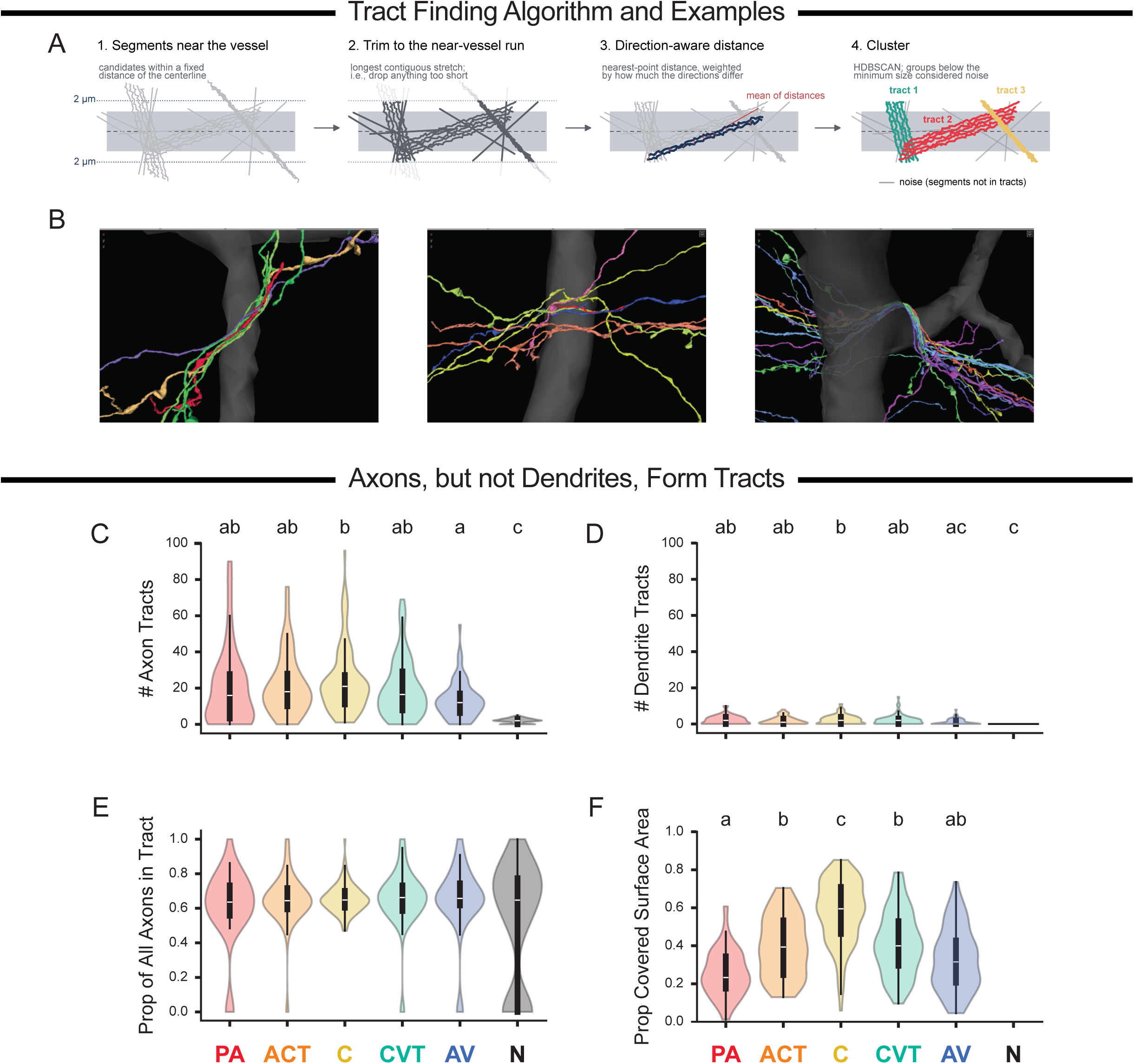
Axons, but not dendrites, form micro-tracts around the vasculature. (A) Steps of the clustering algorithm: we trim axons or dendrites within 2 μm of the vessel to their longest contiguous run in that shell (steps 1 and 2); we compute a direction-aware distance between them (step 3); and we cluster the distance matrix with HDBSCAN (step 4). (B) Three example axonal micro-tracts. These micro-tracts are in https://spelunker.cave-explorer.org/#!middleauth+https://global.daf-apis.com/nglstate/api/v1/6653336237899776, tabs 2 – 4, at positions (i) 198624, 226067, 24174, (ii) 232175, 197731, 21159, (iii) 208512, 220893, 24555. You may have to change the orientation of the view. (C, D) Number of (C) axon and (D) dendrite micro-tracts per branch, by vascular zone; N is the neuropil box. (E) Proportion of the axons near a branch that belong to a micro-tract. (F) Proportion of the branch’s surface area covered by axon micro-tracts; the neuropil has no vessel surface and so is absent. Violins show the distribution, the box the interquartile range and the white line the median. Vascular zones sharing a letter are not significantly different; those sharing no letter differ at adjusted p < 0.05 (Kruskal–Wallis p = 1.03e-29, 9.17e-7 and 1.22e-21 for C, D and F; Dunn’s post-hoc with Bonferroni correction). Letters are not shown in E, where the omnibus was not significant (p = 0.78). n = 34, 41, 108, 80, 70 and 71 branches for PA, ACT, C, CVT, AV and N in panels C and E; 23, 38, 98, 68, 55 and 18 in panel D; 31, 40, 108, 77 and 65 in panel F.

Where micro-tracts form, their internal organization is similar across vascular zones. About 65% of the axons near a vessel join a micro-tract (Figure 3E), which did not differ across vascular zones (Kruskal–Wallis p = 0.78). Axons inside micro-tracts formed synapses at roughly 0.05 to 0.1 per μm (Supplementary Figure 5A), slightly below the rate for axons outside them (roughly 0.07 to 0.11 per μm; Supplementary Figure 5B); this difference is statistically significant but effectively negligible, coming to about three fewer synapses per 100 μm of axon. Within a micro-tract, an axon’s nearest neighbor lay a median of 0.6–0.7 μm away (Supplementary Figure 5C), and neighboring axons diverged by only 7–12° from each other (0° being perfectly parallel; Supplementary Figure 5D). The spacing and alignment of axons within micro-tracts varied modestly across vascular zones (Supplementary Figure 5C, D).

Micro-tracts occupied a median 60% of the capillary surface, 40% of ACT and CVT branches, and 23–32% of PA and AV segments (Dunn’s post-hoc p < 0.05 for capillaries against all other branches; Figure 3F). This ordering is expected from vessel caliber alone, since a micro-tract of a given length or width covers a larger fraction of a thinner vessel, and PAs and AVs tend to have the largest diameter (Supplementary Figure 1B). Thus, of all vessel types, capillaries are most thoroughly covered by axonal micro-tracts.

### Axonal micro-tracts exhibit a range of geometries at the vascular wall

Micro-tracts exhibited diverse orientations around the vessel, with some extending parallel (Figure 4F, vi) or oblique (Figure 4F, i) to the longitudinal axis of the blood vessel. Some micro-tracts barely passed over the blood vessel (Figure 4F, viii), while others wrapped extensively around the circumference (Figure 4F, ix). We developed an algorithm to quantify five aspects of the geometry of micro-tracts in relation to their associated blood vessel (Figure 4A; Methods). We described each micro-tract with five features. Two capture its alignment with the vessel: crossing angle is the mean angle between each axon’s local direction and the vessel’s direction, and vessel fraction is the proportion of the micro-tract’s spatial variance that lies along the vessel axis. One feature captures its reach along the vessel: axial extent is the length of vessel that the micro-tract occupies. Two other features capture how far it wraps around: angular extent is the angle of the vessel’s circumference that the micro-tract spans, and number of turns is how many complete revolutions its axons make. Together, these five features allowed us to quantify the major aspects of micro-tract geometry.

**Figure 4.**
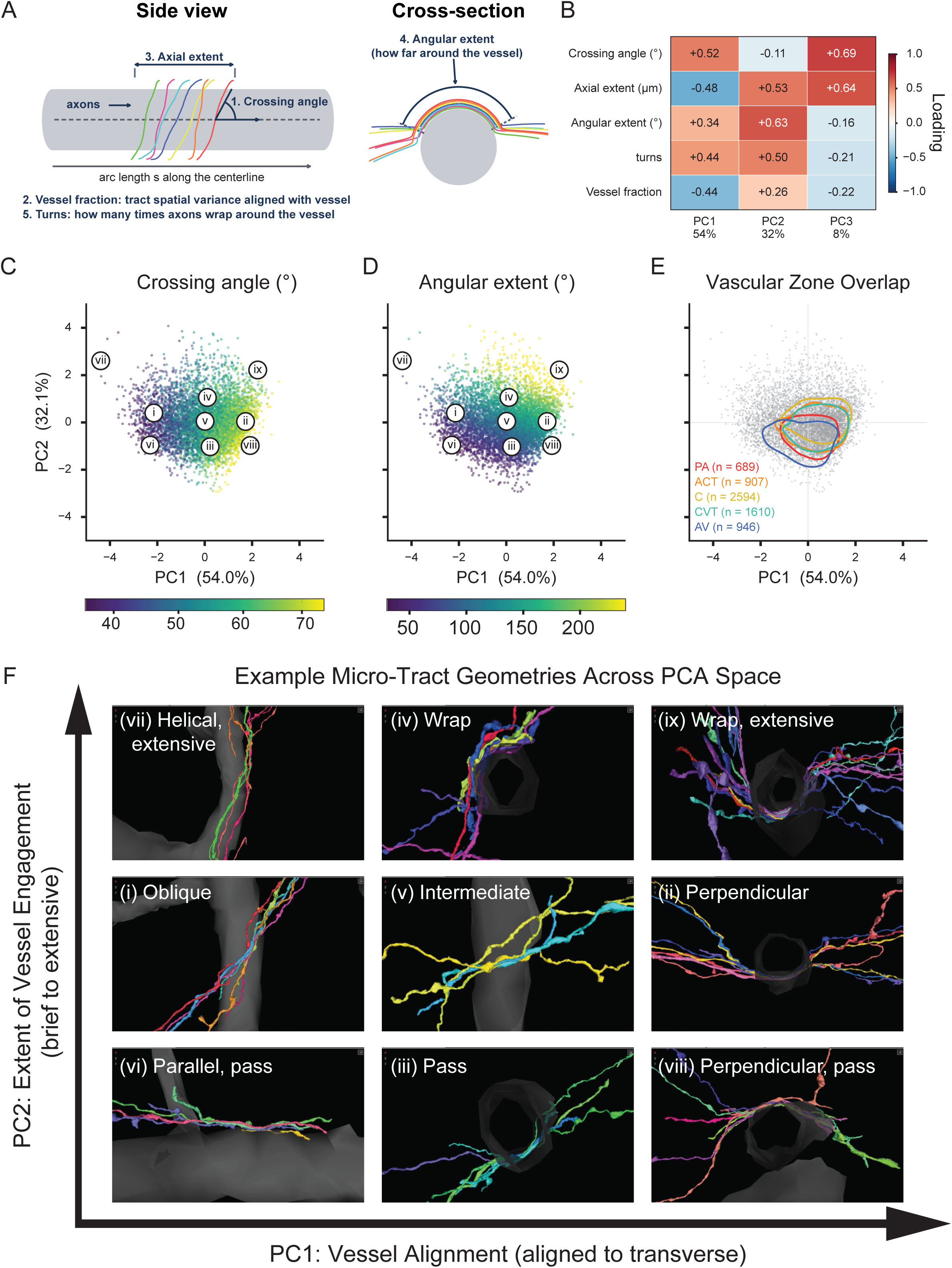
Axon micro-tract geometry varies continuously along two axes — alignment with the vessel and extent of wrapping around it — rather than falling into discrete classes. (A) Schematic of 5 micro-tract geometric features we computed. Colored lines are 7 different axons. (B) PCA loadings for each feature. (C, D) Micro-tracts in PCA space are organized along a spectrum from (C) roughly parallel to perpendicular geometries as their loading on PC1 increases and (D) wrapping more around the circumference of the blood vessel as their loading on PC2 increases. The roman numerals correspond to micro-tracts in F. (E) Micro-tracts with different geometries are found at all vascular zones, though those on PAs and AVs tend to have the smallest loading on PC2 (i.e., they wrap around those vessels less). Contours enclose the densest region containing 50% of each vascular zone’s micro-tracts (kernel density estimate; bandwidth as in Methods). Grey points are individual micro-tracts. (F) Nine example axon micro-tracts, labeled by their geometry and position in PCA space (see panels C and D). These micro-tracts are in https://spelunker.cave-explorer.org/#!middleauth+https://global.daf-apis.com/nglstate/api/v1/5141776185163776, tabs 2 – 10, at positions (i) 198656, 228608, 23808, (ii) 338360, 202790, 24894, (iii) 249516, 181678, 24704, (iv) 197145, 232327, 23647, (v) 119935, 195933, 22729, (vi) 174861, 235201, 22903, (vii) 355699, 170282, 24878, (viii) 347342, 172520, 24885, and (ix) 346041, 203292, 24746.

All five features were unimodally distributed (Supplementary Figure 5E–I) and most pairs were weakly correlated, though four exceeded |Spearman ρ| = 0.5 — the strongest being angular extent with number of turns (ρ = 0.86; Supplementary Figure 6). Even so, no feature was redundant. A micro-tract can reach a high vessel fraction either by running parallel to the vessel or by crossing it perpendicularly while being very wide, so vessel fraction does not determine crossing angle. And because angular extent is bounded at 360°, a micro-tract that wraps three times around the vessel scores the same as one that wraps exactly once; only the number of turns separates them.

To describe this five-dimensional space, we z-scored each feature and applied principal component analysis (Figure 4B). The first two components captured 86% of the variance. On PC1, crossing angle loaded positively (+0.52) while axial extent (−0.48) and vessel fraction (−0.44) loaded negatively. Although no single feature dominated, the pattern of signs describes alignment: a micro-tract with a higher PC1 score meets the vessel more perpendicularly (Figure 4B, C; Supplementary Figure 7A, B). On PC2, every feature except crossing angle loaded positively, led by angular extent (+0.63), axial extent (+0.53) and turns (+0.50). Thus, a higher PC2 score marks a micro-tract that wraps further around the vessel and is generally larger (Figure 4B, D; Supplementary Figure 7C).

To understand what these axes correspond to anatomically, we selected nine micro-tracts spanning the PC1–PC2 plane and rendered them (roman numerals in Figure 4C, D correspond to micro-tracts in Figure 4F). Micro-tracts scoring low on PC1 ran parallel or oblique to the vessel (Figure 4F, i and vi) and those scoring high crossed it perpendicularly (ii, viii, ix); those scoring low on PC2 touched or passed over short segments of the vessel (iii, vi, viii), while those scoring high wrapped around it extensively (iv, vii, ix).

These micro-tracts did not fall into distinct geometric classes. Instead, their geometry is continuous: no feature was bimodal (Supplementary Figure 5E–I), no feature pair separated into distinct groups (Supplementary Figure 6), and the PC1–PC2 plane held a single connected cloud (Figure 4C, D). The intermediate example (Figure 4F, v) is the case in point: aligned obliquely rather than parallel or perpendicular, and wrapping the vessel a little but not much.

Finally, micro-tracts of different geometries were not concentrated on any one vessel class (Figure 4E), except that those on PA and AV segments scored lower on PC2, perhaps because these are the largest caliber vascular zones we analyzed (median diameters of ∼7 μm versus 4–5.5 μm for the rest; Supplementary Figure 1B). Indeed, across all 6,745 micro-tracts, PC2 declined with the diameter of the host vessel (Spearman ρ = −0.50), as expected if a wider vessel is harder to wrap around.

### The axons of basket cells approach the most capillaries

The cerebral cortex contains several different neuron subtypes, each of which may have a unique role in NVC. Parvalbumin-positive (PV+) interneurons, for instance, have been shown to regulate NVC in a depth-dependent manner (Rakymzhan et al., 2025). PV+ interneurons in the mouse visual cortex also integrate activity from excitatory neurons over a ∼75 μm-wide region (Scholl et al., 2015), which suggests their activity levels may be a good readout of local network activity. This suggests that PV+ interneurons may contribute to NVC if their axons course near blood vessels. It was not possible to determine which neurons form micro-tracts in the MICrONS dataset, since not all axons are proofread or connected to somata. However, it was possible to ask a related question: of the proofread neurons, how often did their axons come close to any blood vessel?

We examined 2,144 proofread neurons, whose axons and dendrites are complete and validated, and computed (i) the minimum distance between the neuron’s processes and any blood vessel (Figure 5A), and (ii) the number of distinct vessel branches that came within 2 μm of them (Figure 5C). At their closest, the axons of every cell type came within a median of 0.2–0.3 μm of a vessel (Figure 5B), and dendrites within 0.2–0.4 μm (Supplementary Figure 8A). Somata (Supplementary Figure 8B) and axon initial segments (AIS; Supplementary Figure 8C), on the other hand, remained a median of 10–13 μm away. Thus, by this metric, every cell type’s axon or dendrite has the potential to “innervate” a blood vessel, or join a micro-tract.

**Figure 5.**
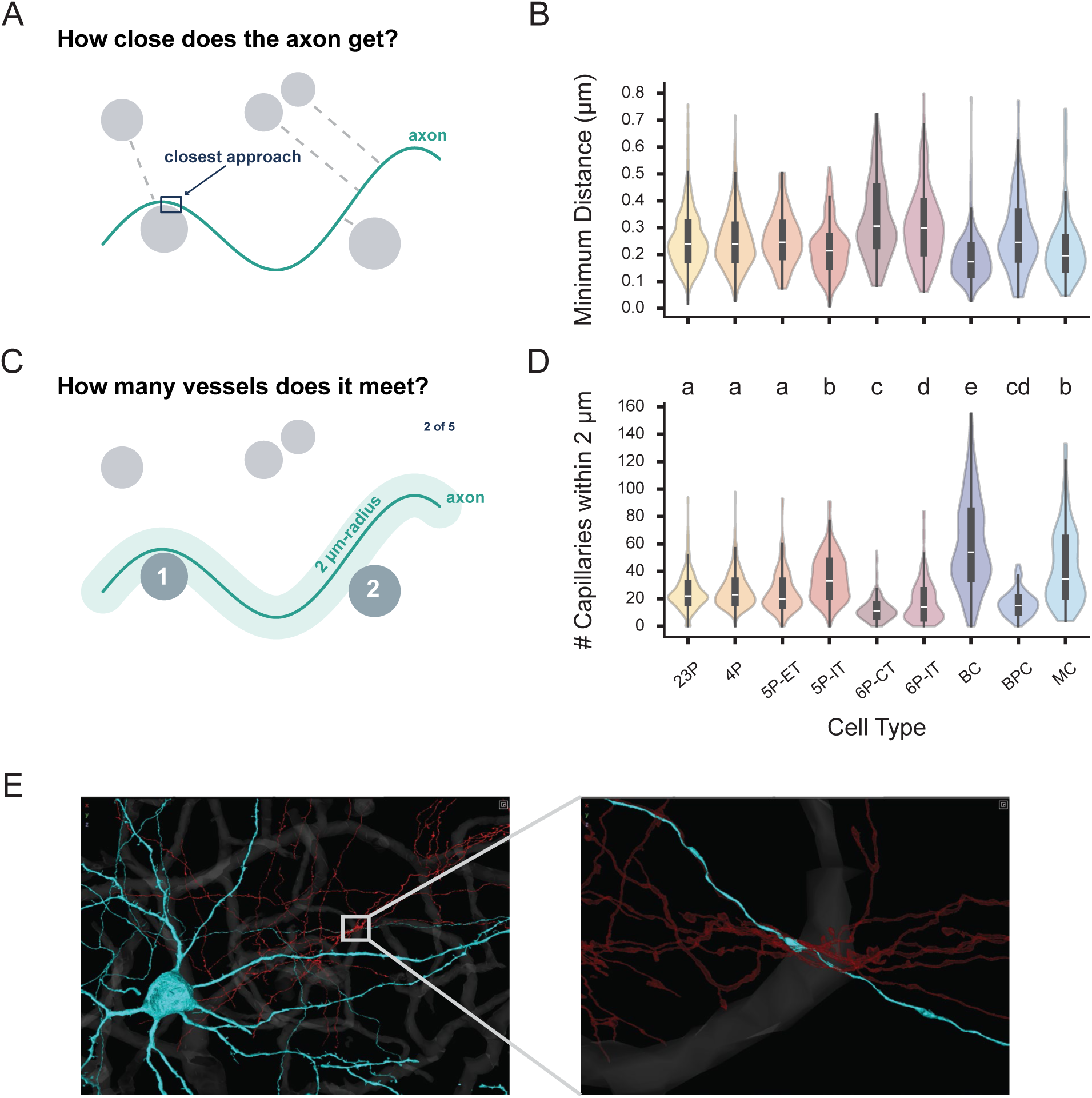
Basket cell axons are candidate members of micro-tracts. (A) Schematic 1: how close does a cell’s axon get to any blood vessel? (B) Answer to Schematic 1: The axons of all cell types get similarly close to blood vessels. (C) Schematic 2: how many different blood vessel branches lie within 2 μm of a cell’s axon? (D) Answer to Schematic 2: BC axons get within 2 μm of more capillaries than other cell types. (E) Example of a Basket cell’s (BC) axon (cyan) in a micro-tract (axons from other cells are red; https://spelunker.cave-explorer.org/#!middleauth+https://global.daf-apis.com/nglstate/api/v1/4817557316435968). Inset shows the site of contact with the blood vessel (grey); coordinates are 198494, 226571, 25284. Violins show the distribution, the box the interquartile range and the white line the median. In D, cell types sharing a letter are not significantly different; those sharing no letter differ at adjusted p < 0.05 (Kruskal–Wallis p = 1.8e-127; Dunn’s post-hoc with Bonferroni correction). Differences in B were also detectable (Kruskal–Wallis p = 5.0e-42), but every cell type’s median lies between 0.2 and 0.3 μm, so letters would overstate them and are not displayed.

We next asked how many vessel branches lie within 2 μm of each cell type’s processes (Figure 5C). Basket cell (BC) axons – the PV+ morphological subtype – came within 2 μm of a median of 54 capillaries, more than any other cell type (Dunn’s post-hoc p < 0.05 for every cell type against BC; Figure 5D, Supplementary Table 1); Figure 5E shows an example of a BC’s axon joining a micro-tract running close to a capillary. Martinotti cells (MCs) and 5P-IT cells followed, at medians of 34 and 33, respectively, each differing from every other cell type but not from one another; the remaining cell types reached only 11–23 capillaries.

The dendrites gave a different ranking. There, 5P-ET (extratelencephalic-projecting) neurons – an L5 excitatory subtype projecting both within the telencephalon and to subcerebral regions – came within 2 μm of a median of 29 capillaries, compared to 8–17 for every other type (Dunn’s post-hoc p < 0.05 for every cell type against 5P-ET; Supplementary Table 1). Neither BC nor 5P-ET cells stood out in the other compartment: BC dendrites reached a median of 17 capillaries and 5P-ET axons a median of 20, both indistinguishable from several pyramidal classes.

Contacts with the other vascular zones were far rarer (median 0–2; Supplementary Table 1), as expected given how many more capillary branches the volume contains; the informative comparison is between cell types within capillaries, as above.

Why are BC axons and 5P-ET dendrites near so many capillaries? Each has an unusually long arbor in the compartment that stands out: BC axons are among the longest of any cell type (median 16.1 mm; Supplementary Figure 9A) while their dendrites are unremarkable, and 5P-ET dendrites are the longest (median 7.2 mm; Supplementary Figure 9B) while their axons are unremarkable. Normalizing the number of nearby capillaries by the total length of each process removes the difference: across every cell type and both compartments, rates fell between 0.0034 and 0.0045 capillaries per μm, with BC and MC at the low end rather than the high (Supplementary Figure 9C, D). The number of capillaries a neuron approaches is therefore set by how much arbor it has, not by a subtype-specific tendency to seek vessels out.

## Discussion

The chemical signals involved in neurovascular coupling (NVC) have been extensively studied, but the anatomical substrate that supports it is less understood. To this end, we systematically characterized the cellular milieu around the vasculature in a ∼1 mm^3^ volume of mouse visual cortex acquired using electron microscopy (MICrONS Consortium, 2025). We found that neuronal components, such as axons, dendrites, and synapses, occupy a larger proportion of the volume near capillaries, capillary-venule transition zones (CVTs), and ascending venules (AVs), than near penetrating arterioles (PAs) and arteriole-capillary transition zones (ACTs). Since signals released from active neurons trigger NVC (Iadecola, 2017; Longden et al., 2017), this suggests that capillaries and the venous vessels (CVTs and AVs) are exposed to a higher concentration of those signals than the arterial side of the vasculature. This largely agrees with a model in which capillaries, which course through the parenchyma and are within ∼13 μm from most neurons, are ideally situated to sense local neuronal activity. Our findings reveal a structural framework that could support capillary-to-arteriole signaling demonstrated by Longden et al. (2017), where K^+^ release from active neurons hyperpolarize capillary endothelial cells, which spreads retrogradely towards the PAs to their dilation. Thus, capillary networks are emerging as an important player in NVC, and our work supports this model.

### Axons organize into micro-tracts that are enriched at capillaries

We found that axons near blood vessels organize into ∼20 micro-tracts per branch segment and, though axons occupy a larger proportion of the volume in the parenchyma than near blood vessels, they rarely form micro-tracts there (Figure 3). Furthermore, though all vascular zones have a similar number of micro-tracts, these micro-tracts cover a larger proportion of the surface area of capillaries, likely due to their smaller diameter. There are two questions: what is the role of micro-tracts in NVC, and how do they form?

We suggest that micro-tracts can induce vasodilation or vasoconstriction depending on their location. As action potentials propagate down an axon, they regenerate by importing sodium (Na^+^) and exporting K^+^. When unmyelinated axons bundle into micro-tracts, modeling work has shown that they can depolarize each other by ∼29 mV for a 100 mV action potential via ephaptic coupling (Bokil et al., 2001). Thus, axons in a micro-tract may trigger action potentials in each other, which would enable the expelled K^+^ to reach a higher concentration than if the axons were diffusely distributed. At capillaries, then, micro-tracts could facilitate the hyperpolarization of endothelial cells, which would dilate upstream PAs and increase local CBF (Longden et al., 2017). However, micro-tracts may also be involved in vasoconstriction. Capillary-mediated vasodilation occurs at 10 mM K^+^ but fails at concentrations of 25 mM or above (Longden et al., 2017). Furthermore, extracellular K^+^ less than 20 mM dilates PAs, while higher concentrations induce constriction (Girouard et al., 2010). Altogether, at PAs, the number of co-active axons and micro-tracts could induce either vasodilation or vasoconstriction, while at capillaries they would trigger vasodilation. Future work will investigate whether axons in micro-tracts are ever co-active, under what conditions, and if the micro-tracts at PAs can switch between dilation and constriction.

We propose that axons typically form micro-tracts near blood vessels because astrocytes build a scaffold in the form of the extracellular matrix (ECM). Astrocytes line most of the vasculature (Diaz-Castro et al., 2023), with few gaps (Zhang et al., 2024), and they secrete a host of carbohydrates and proteins that act as a scaffold for the intercellular milieu (Ge & Tajerian, 2026); this mixture is called the ECM. Some components of the ECM, such as laminin, play a role in axon growth and guidance (Pires Neto et al., 1999). More generally, axons can reach distant locations during development via haptotaxis – directed movement of cells along surfaces bound with an adhesive molecule (Varadarajan et al., 2017). Thus, astrocytes might encourage the formation of axon micro-tracts during development via the formation of the ECM.

We also found that these micro-tracts exhibit a continuum of morphologies along two major geometric axes: their alignment with the blood vessel (some run parallel, others perpendicular) and how much they wrap around the blood vessel (between ∼40° and ∼210° of the vessel circumference). Future work should establish which geometry would be most effective at increasing the local concentration of K^+^ near capillaries and PAs.

### Basket and 5P-ET cells contact many capillaries

Finally, we found that the axons of basket cells (BCs) approach a median of 54 capillaries. This suggests that BCs might be common members of micro-tracts, and that they may exert their role by hyperpolarizing capillary endothelial cells to induce vasodilation. Furthermore, BCs can fire at up to 582 Hz (Wang et al., 2016) and integrate neuronal activity over a range of ∼100 μm in mouse visual cortex (Scholl et al., 2015). Therefore, BCs are well-suited to monitor local neuronal activity and to convey this information to capillaries by dumping K^+^ at very high rates, which would increase the local K^+^ concentration and ultimately trigger upstream vasodilation and an increase in CBF that supports ongoing activity. Future work will look for all micro-tracts in this EM volume and comprehensively assess which cell types most often join micro-tracts, and in which combinations.

We also found that the dendrites of 5P-ET neurons reach a median of 29 capillaries, more than all other cell types. 5P-ET neurons are considered the main outputs of the cortical microcircuit: they integrate inputs from throughout the cortical layers (they have extensive basal and oblique dendrites and a thick apical dendrite that reaches L1) and their axon reaches far (they have a sparse local axonal arbor and long-range cortical and subcortical projections) (Bodor et al., 2025; Ramaswamy & Markram, 2015). Moreover, their primary local synaptic partners are inhibitory interneurons, such as BCs and Martinotti cells, where the former innervate near the soma of 5P-ETs and the latter innervate their distal dendrites and terminal tufts (Bodor et al., 2025; Ramaswamy & Markram, 2015). This implies that there may be an interplay between neurons whose axons contact capillaries at high rates (BCs) and neurons whose dendrites do the same (5P-ET neurons). While action potentials propagating along dendrites will also result in K^+^ efflux, it is unclear how the complex distribution of voltage-gated ion channels along the dendrites of 5P-ET neurons may affect nearby blood vessels (Ramaswamy & Markram, 2015). Future work should establish how often the dendrites of all cell types, particularly 5P-ET neurons, contact blood vessels; if these contacts are specific to particular vascular zones; which dendritic region (i.e., basal, oblique, apical, or terminal), if any, contacts blood vessels; and how BCs and 5P-ET neurons differentially or synergistically mediate NVC.

More generally, we found that, normalized by total length, all cells reached similar numbers of capillaries. Cortical energy use is dominated by action potentials (Attwell & Laughlin, 2001; Harris et al., 2012), which propagate through axons and dendrites alike. Thus, any cell with a larger axonal or dendritic arbor will need more oxygen and glucose, which means it will need to be closer to more blood vessels. Simply put, this shows that (blood) supply tracks (energetic) demand.

## Methods

### Datasets

The electron microscopy (EM) data came from the MICrONS project (MICrONS Consortium, 2025). Briefly, a 1.3 × 0.87 × 0.82 mm^3^ region of visual cortex of a 87-day-old male mouse was reconstructed using serial section transmission EM at a resolution of 8 × 8 × 40 nm^3^. The resulting imagery was segmented and underwent various rounds of proofreading; we used version 1078, released June 2024, for all analyses except for downloading proofread cells, for which we used version 1621. The blood vessels were previously segmented at a coarser resolution of 320 × 256 × 256 nm by Wan et al. (2025), which we downloaded at 1280 × 1024 × 1024 nm.

### Data Analysis

#### Vascular Segmentation Correction, Skeletonization, and Branch Characterization

We converted the 3D vascular segmentation to a 3D skeleton, or single-voxel wide line that traces a path through the center of all blood vessels. We first found the largest connected component (skimage.measure.label) and filled its internal holes and cavities with an iterative neighbor-voting rule (Simple ITK’s VotingBinaryIterativeHoleFilling using default settings). We then converted the segmentation to a skeleton using using Lee’s algorithm (skimage.morphology.skeletonize) (Lee et al., 1994); however, the skeleton still had two main problems: 1) bristles (short terminal edges, usually on large branches) and 2) short cycles (topological paths that formed a loop, usually on large branches). To address these problems, we iterated between fixing both:

1. We removed bristles that were less than 35 microns long and not touching the boundary of the imaged volume (networkx).
2. We removed all cycles that are less than 1000 microns long (effectively, all cycles) and isolated from the rest of the skeleton (skan).
3. We removed bristles that were less than 30 microns long.
4. We randomly broke 1 edge in each triangle (networkx).
5. We iteratively condensed cycles that were less than 50 microns long. We stopped when the number of cycles did not change or we had iterated through the loop 10 times. Throughout, we reconnected branches that were less than 20 microns away along the skeleton prior to any modifications (i.e., before step 1) by simply copying the old path to the new skeleton, and we removed isolated parts of the skeleton (components) that were less than 2 microns long.
6. We removed bristles that were less than 2 microns long.
7. We reconnected branches as in Step 5.
8. We merged high-degree nodes that were less than 3 microns away from each other by removing both and adding a new node at their spatial centroid, and we connected their old partners to the new node; we repeated this step until the number of nodes in the skeleton did not change.
9. Throughout Steps 1 through 8, we periodically converted the skeleton to a graph (networkx) and removed self-loop edges.

This procedure generated a skeleton for the largest connected component of the vascular network with several desirable properties: no errant cycles, no errant bristles, and connected into a single component. We then used VesselExpress to find and morphologically characterize all branches (i.e., sections of the skeleton between two branch points; n = 11380 branches) of the vascular network (Spangenberg et al., 2023). We then manually removed branches that were merges of two or more branches due to problems with the segmentation or issues at the boundary of the imaged volume, for a total of n = 11345 branches.

The features quantified by VesselExpress for each branch *s* are described in (Spangenberg et al., 2023), and are summarized here: 1) branch length (path distance along the skeleton between two branch points, in microns), 2) mean diameter (mean of the Euclidean distance transform evaluated at each node in the branch skeleton, in microns), 3) tortuosity (one minus the Euclidean distance between the branch points divided by the branch length), and 4) volume (using the standard formula for a cylinder; Equation 1).

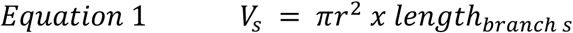

We then computed the surface area of each branch *s* using the standard formula for a cylinder (Equation 2).

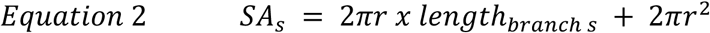

#### Vascular Zone Annotation

We used the Connectome Annotation Versioning Engine (Dorkenwald et al., 2025) point annotation tool to manually label branches into one of ten categories: pial (P), penetrating arteriole (PA), ascending venule (AV), principal cortical vein (PCV) arteriole-capillary transition zone (ACT), capillary-venule transition zone (CVT), deep CVT (branching off of PCVs), terminal PA, terminal AV, and terminal PCV.

##### Major vessels

P branches were identified as the large diameter vessels running horizontally along the surface of the brain. PAs and AVs were initially identified as large-diameter branches penetrating from the pial surface straight down into the brain. Branches were annotated as PA when encircled by ring-like smooth muscle cells (Hartmann et al., 2022), or AV if they lacked these cells. PCVs were identified as two large-diameter branches running horizontally along the white matter (the imaged volume did not have the vertical component penetrating across layers of cortex), and we labeled their full extent using a previously described procedure (Stamenkovic et al., 2025). Briefly, we first found the capillary network for each PCV branch, defined as every capillary branch that converges flow toward it. We defined the upstream boundary as the last branch that clearly converged to the PCV and not an AV. We then traced a path towards the PCV, and labeled the first branch ≥ 10 μm in diameter, and everything downstream, as the PCV.

##### Transition vessels

ACTs, CVTs, and deep CVTs were identified as the first order branches off of the main vessel. If the first order branch was particularly small (< ∼5 microns), we annotated the immediately downstream branches as transition vessels, too. These transition vessels often emanated orthogonally to the direction of the major vessels.

##### Terminal vessels

Terminal zones for the major vessels were identified as the region at which the major vessel split, usually in a Y-shape, into two or more daughter vessels with similar diameters.

After finishing the initial annotations, we iteratively looked for annotation errors (e.g., branches between two AVs should be an AV, but may have been missed; or capillaries attached to a PA should be an ACT but may have not been labeled as such) until we detected no more errors.

This link – https://spelunker.cave-explorer.org/#!middleauth+https://global.daf-apis.com/nglstate/api/v1/5249453187923968 – will allow anyone to explore the dataset.

#### Capillary Network Annotation

We represented the vascular network as a skeleton graph (nodes are 3D skeleton voxels and edges connect adjacent voxels), branch graph (nodes are branches and edges connect branches with a common branch point, or junction), and line graph (nodes are branch points and edges are branches); the latter two are more efficient for computing certain properties, and the line graph in particular allowed us to use branch properties (e.g., length) as edge weights.

We annotated each skeleton node with its branch identity, vessel type, radius, length, straightness, branching angle, and distance to the outer surface of the vasculature. We assigned junction nodes, which belong to no single branch, to the most common non-capillary type among their neighbors, or Capillary if all their neighbors were capillaries.

To restrict analysis to the capillary bed, we removed branches classified as PAs, AVs, PCVs, or Ps, and kept the largest connected component of the remainder; thus, every remaining C (capillary) could only reach a blood source or drain by going through the rest of the capillaries. We defined the ACT and terminal PA branches to be the blood sources, and the CVT, terminal AV, deep CVT, and terminal PCV branches as the blood drains. We removed all branches that were not directly connected to at least one capillary, and capillaries connected to a marge error (see **Vascular Segmentation Correction, Skeletonization, and Branch Characterization**). Thus, though our capillary network had n = 9,292 capillaries, we only processed n = 7,757 capillaries as follows.

We computed the shortest paths between all sources and drains on the line graph, both unweighted (branch count) and weighted (branch length), using Dijkstra’s algorithm (rustworkx.dijkstra_shortest_paths). We assigned each capillary branch a branch order and geodesic distance (distance along the vascular network) relative to the nearest source and drain.

#### Gini Coefficient

We found all shortest paths – not weighted by length – between each pair of blood sources (i.e., ACT, terminal PA) and drains (i.e., CVT, deep CVT, terminal AV, terminal PCV), and counted how many passed through each processed capillary. Of n = 7757 capillaries, 3.8% lay on no shortest path. Let *v*₁ ≤ *v*₂ ≤ ⋯ ≤ *vₙ* denote these traversal counts per capillary in ascending order. The Lorenz curve plots the cumulative share of capillaries, *i*/*n*, against the cumulative share of traversals, (Σ*_j_*_≤*i*_*v_j_*)/(Σ*_j_v_j_*), so that the diagonal corresponds to every capillary carrying equal traffic. Note that the denominator Σ_j_*v*_j_ is not the number of paths: a path contributes one traversal to each capillary it crosses, so a path crossing *k* capillaries contributes *k*.

The Gini coefficient was computed directly from the sorted counts as

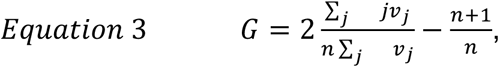

which is equal to twice the area between the Lorenz curve and the diagonal. *G* = 0 when all capillaries carry identical traffic and attains its maximum of (*n* − 1)/*n* when a single capillary carries all of it. The share carried by the busiest decile was taken as one minus the cumulative share of the least busy 90% of capillaries.

#### Segment Classification

We chose n = 347 branches that were PA (n = 38), ACT (n = 41), C (n = 110), CVT (n = 80), or AV (n = 78), spread across all layers of cortex, and n = 80 Neuropil regions – 20 per layers 2/3, 4, 5, and 6. All n = 427 regions were in minnie65, where the EM segmentation exists, and at least 100 voxels in size. For each branch, we downloaded segments that were up to 5 μms away from the surface of the branch; we dropped voxels that were equidistant to 2 or more different branches. For each Neuropil region, we found a centroid that was approximately 13 μms away from the nearest blood vessel – the median distance between any Neuropil voxel and any blood vessel (Supplementary Figure 1E) – and downloaded a 5 x 5 x 5 μm region around it.

To classify segmented objects within these volumes, we trained a logistic regression classifier on SegCLR embedding vectors (Dorkenwald et al., 2023), which were previously shown to be capable of accurately discriminating between broad categories of segmented objects in connectomics volumes (e.g., axon vs. dendrite vs. glia). Briefly, SegCLR applies a convolutional neural network to the EM imagery at a given point, masked by the segmentation for that object. The network is trained with contrastive learning (Chen et al., 2020) to discriminate nearby points on the same segmented object from random other objects sampled from the volume. Once the network is trained, the “embedding” associated with SegCLR is the *d*-dimensional readout from hidden units in the network for a given segmentation/imagery cutout for that object (here, d = 64).

We leveraged these embeddings (previously computed for MICrONS at version 943) to train a classifier to label objects as being axon, dendrite, soma, glia, large axons (e.g. myelinated), or a broad perivascular class. Labels for the axon, dendrite, soma, and glia classes came from manually proofread local cells (CAVE table “proofreading_status_and_strategy”, version 1098). We augmented this dataset with additional labels for large axons (often myelinated) which came from outside of the volume, as well as objects directly contacting or surrounding the vasculature (the perivascular category). This latter category largely comprises endothelial and mural cells (e.g., smooth muscle cells, pericytes). We mapped points on the skeletons for these objects to their corresponding SegCLR embeddings as their vector representations, and then trained a logistic regression classifier to predict labels from these features (Pedregosa et al., 2011). The classifier used default scikit-learn parameters besides max_iter=2000 and class_weight=”balanced” to more equally weight the importance of points from each category.

#### Micro-tract Detection

After classifying segments in each region of interest (ROI), we detected micro-tracts. We first extracted the 3D voxels corresponding to segments that fulfilled various criteria: 1) classified as axons with a posterior probability ≥ 0.75, 2) size greater than a threshold based on myelinated axons (∼4e7 nm^3^; Supplementary Figure 3A), 3) associated root IDs were most often classified as axon, 4) distance between their endpoints was at least 5 μms (which excludes both short fragments and highly tortuous segments that lack a consistent direction), and 5) length of the axon contained with 2 μms of the blood vessel was at least 5 μms. We then kept the longest section of each axon within 2 μms of the blood vessel.

We then computed a pairwise direction-aware distance between each pair of axons. For each point on one axon, we found the nearest point on the other and multiplied that distance by (1 + α·sin θ), where θ is the angle between the two axons’ local directions at those points and α = 2.0; local direction was taken as a centered difference over ±5 points (∼2.6 μm). We then averaged the distance from axon A to B and B to A to generate a symmetric *n x n* distance matrix *D*. We then used HDBSCAN* – a hierarchical clustering algorithm that clusters dense areas and calls the rest noise (Campello et al., 2013) – to cluster *D* into clusters with at least 5 axons; each cluster is a micro-tract.

#### Micro-tract Geometry

We described each micro-tract in a coordinate frame centered on the blood vessel branch with which it associates. We converted coordinates for each branch and axon from voxels to microns. We fit the branch’s skeleton with a cubic spline, uniformly resampled it, and then propagated a parallel-transport (Bishop) frame along it (Bishop, 1975). We assigned every axon coordinate to its nearest spline coordinate, which allowed us to derive, per axon coordinate, its arc length *s* along the vessel and angle *θ* around the vessel.

We computed five features per tract:

1. **crossing angle degree**: the mean acute angle between each axon’s local direction and the vessel’s direction at the nearest centerline point, where 0° indicates running along the vessel (parallel) and 90° crossing it (perpendicular). Local direction was estimated from a windowed tangent spanning roughly *N*/15 points since point-to-point differencing is dominated by coordinate noise at this sampling density. The micro-tract-level property is the median over axons.
2. **vessel fraction**: the proportion of the tract’s total spatial variance lying along the vessel axis. This measures the same parallel-versus-perpendicular contrast as crossing angle degree, albeit at the level of the whole point cloud rather than for individual axons. Vessel fraction and crossing angle degree compensate for each other; for instance, a broad sheet whose axial extent exceeds its arc length has an elongated point cloud and reads as parallel by this measure, but the crossing angle degree correctly identifies its individual axons as perpendicular.
3. **axial extent**: the range of *s* spanned by the tract, in μms.
4. **angular extent:** the extent of the vessel circumference spanned by the tract’s voxels, in degrees. We divided the circumference into 10° bins, assigned each axon to the bins its voxels occupied, and kept bins with at least two distinct axons; the extent is the smallest arc containing those bins. Note that this can underestimate the angular extent, mostly because axons can be fragmented into different segments, thus making it hard for any one bin to reach the four-axon threshold.
5. **number of turns:** median of how many times each axon turns around the vessel.

#### Distance Between Cells and Vasculature

We downloaded 2,191 cells with proofread axons and dendrites from table aibs_metamodel_celltypes_v661, version 1621, and kept cell types with at least 30 cells; thus, we dropped the 5P-NP and neurogliaform cells, for a final n = 2,155 cells. We downloaded the mesh of the vascular segmentation (Wan et al., 2025). We computed skeletons for each neuron using pcg_skel (https://github.com/CAVEconnectome/pcg_skel/) which implements the TEASAR algorithm on a coarse graph of the segmentation (Sato et al., 2000). From these skeletons, we used ossify (https://github.com/ceesem/ossify) to classify skeleton segments as axon, dendrite, or soma, with the soma being defined as the root of the skeleton anchored to the location of the detected nucleus of the cell, and using the “label_axon_from_spectral_split” function in ossify to label axon and dendrite. This method treats input synapses on the skeleton as “dendrite” and output synapses as “axon”, and applies a smoothing approach to extend these labels to the rest of the skeleton nodes. For each cell, we masked out the axon, dendrite, or axon initial segment (AIS) (https://github.com/AllenInstitute/em_skeleton_feature_extraction), and found its distance to the nearest blood vessel mesh. The AIS was defined as the part of the axon that is closest to the soma, where distance was computed on each neuron’s skeleton; the axon and dendrite regions were previously identified. For a subset of n = 496 neurons, we then computationally located all points along their axons that were within 2 μm of any blood vessel, and tagged those points as: 1) “contact” between axon and blood vessel, 2) “intervening astrocyte” when astrocytic endfeet separated the axon and blood vessel, and 3) “unclear” when we could not distinguish between the two possibilities due to issues with the resolution and segmentation.

#### AI Disclosure

We used Claude (Anthropic PBC; Opus 4.6 and 5) to generate code and draft parts of the Methods section. We manually verified all outputs.

---

**Supplementary Figure 1.**
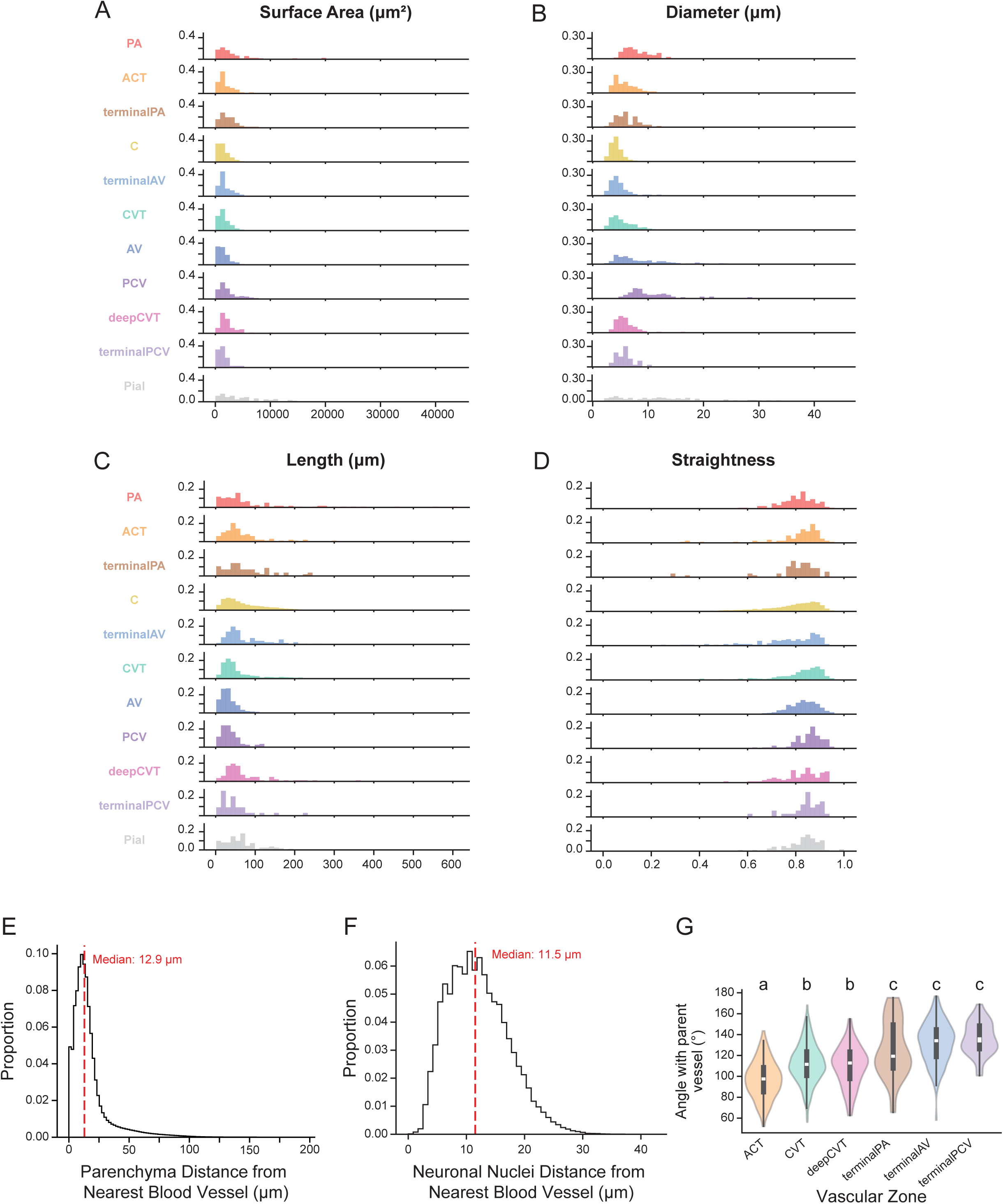
Vasculature morphology. Distributions of (A) surface area, (B) diameter, (C) length, and (D) straightness of each vascular zone. (E) Distribution of distances between neuropil and nearest blood vessel. (F) Distribution of distances between neuronal nuclei and nearest blood vessel. (G) Angle each vascular zone makes from its parent branch. Violins show the distribution, the box the interquartile range and the white line the median. Vascular zones sharing a letter are not significantly different; those sharing no letter differ at adjusted p < 0.05 (Kruskal–Wallis p-value 1.61e-37; Dunn’s post-hoc with Bonferroni correction).

**Supplementary Figure 2.**
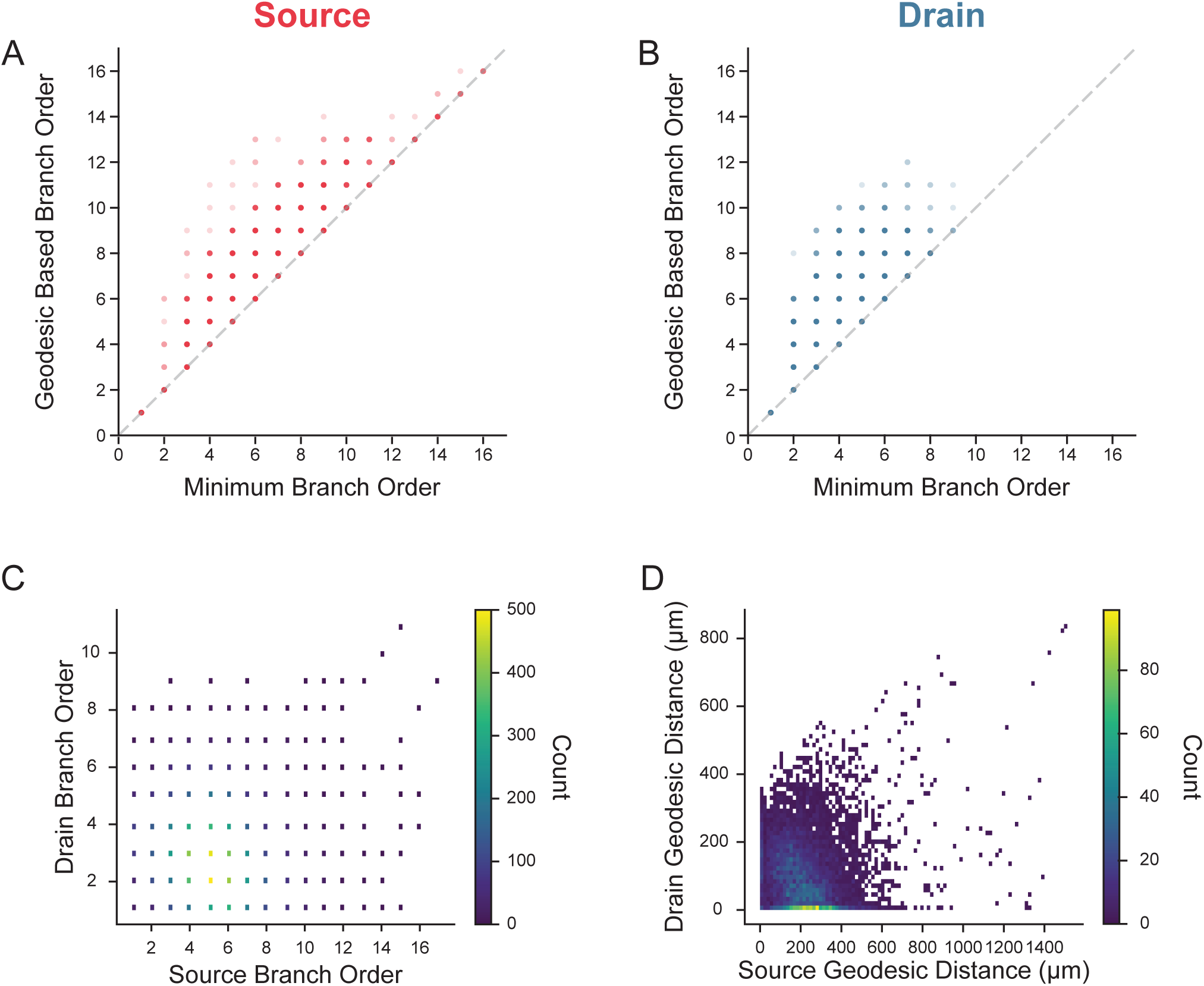
Branch order differs between geodesic-distance-based and minimum-branch-order-based definitions. (A, B) Number of branches along the geodesic route (y-axis) plotted against the minimum branch order (x-axis), for each capillary’s nearest (A) source and (B) drain. Points on the dashed identity line are capillaries for which the two definitions agree; points above it are those whose geodesic route passes through additional branches. Color saturation reflects the number of overlapping capillaries. (C, D) Joint distribution of each capillary’s distance to its nearest source and to its nearest drain, expressed as (C) branch order and (D) geodesic distance; color gives the number of capillaries per bin. n = 7,757 capillaries.

**Supplementary Figure 3.**
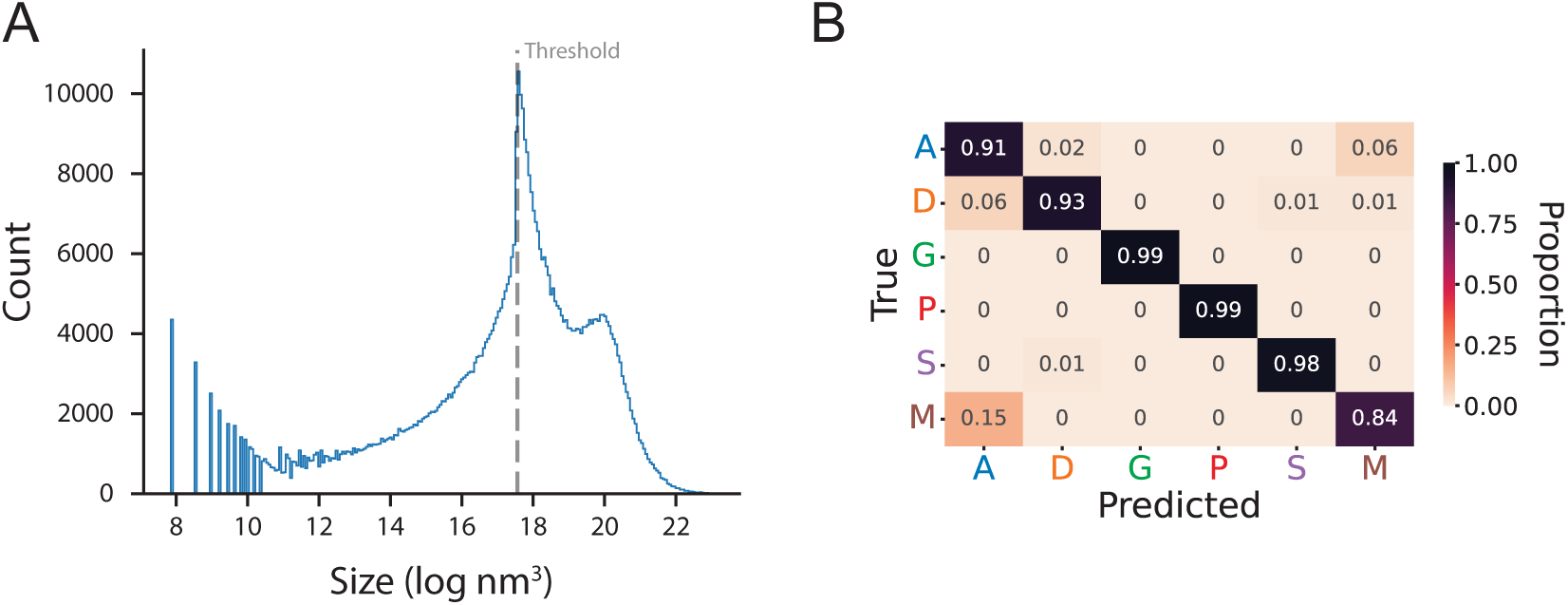
Segment classification. (A) Histogram of segment sizes. Segments below the threshold of ∼4e7 nm^3^ were not analyzed. (B) Confusion matrix shows our classification is accurate. A: axon, D: dendrite, G: glia, P: perivascular, S: soma, and M: thick/myelinated axon.

**Supplementary Figure 4.**
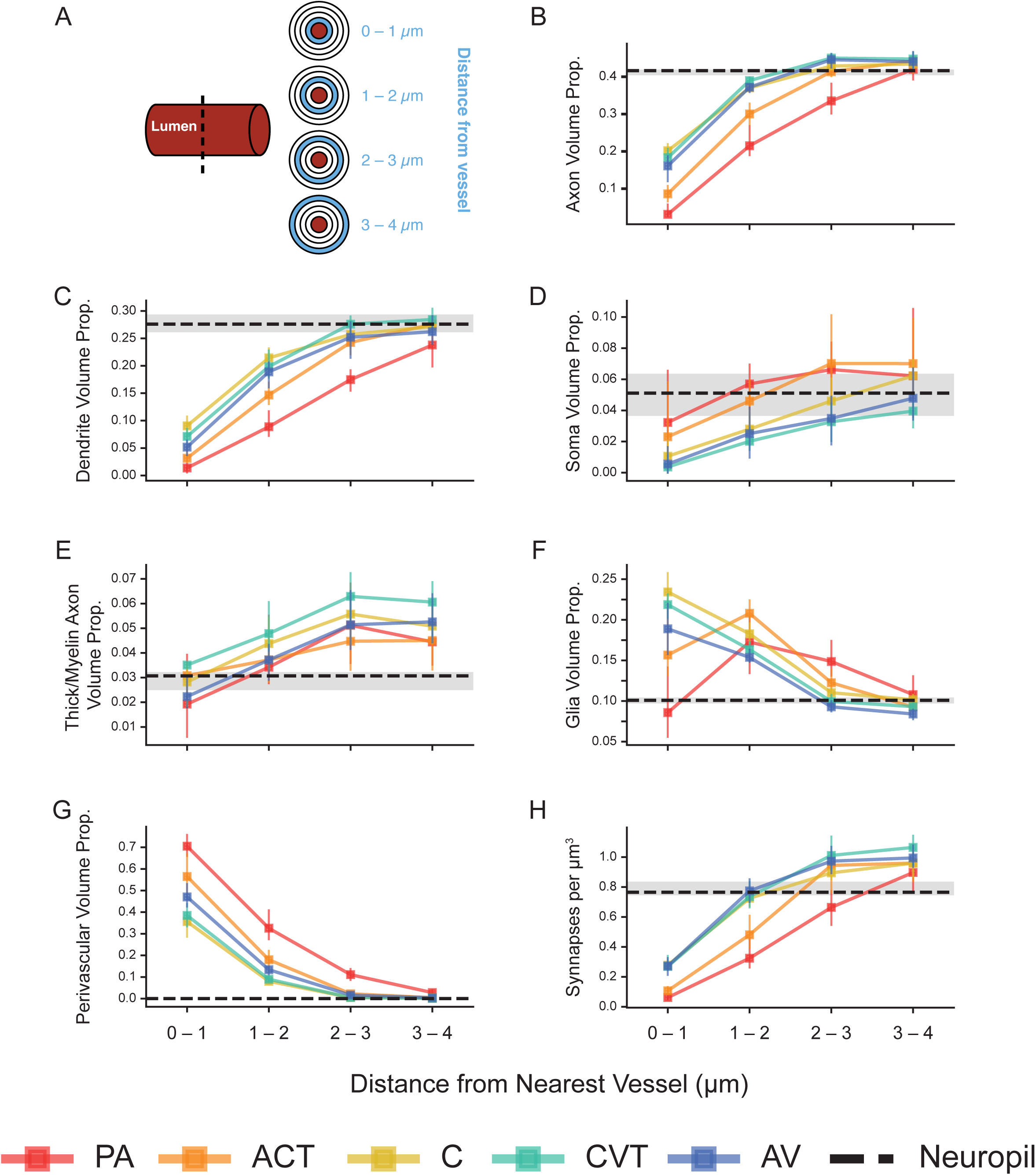
Composition of the neighborhood around blood vessels. (A) Volumes were measured in concentric 1 μm shells around each vessel’s lumen. (B – G) Volume proportion of each compartment as a function of shell: (B) axons, (C) dendrites, (D) somata, (E) thick and myelinated axons, (F) glia, and (G) perivascular tissue. (H) Synapse density as a function of shell. Squares are medians and error bars are 95% bootstrap confidence intervals across branches; the black dashed lines and shaded bands give the median and 95% bootstrap confidence intervals for the Neuropil boxes.

**Supplementary Figure 5.**
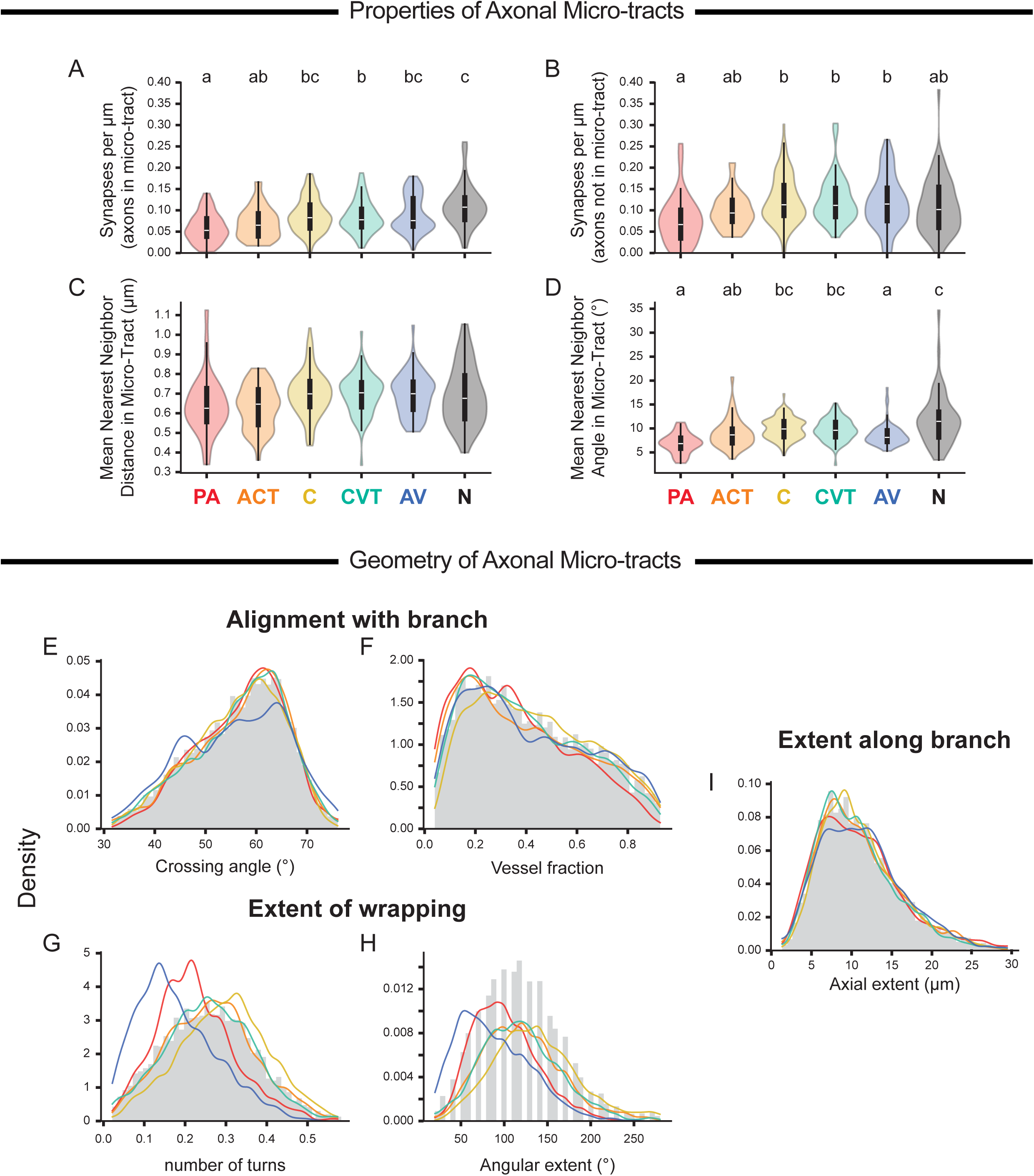
Micro-tracts are similar across vascular zones. (A, B) Number of synapses per μm for axons (A) in micro-tracts and (B) not in micro-tracts for each vascular zone. (C, D) Coherence of axon micro-tracts: (C) mean nearest neighbor distance and (D) mean nearest neighbor angle per axon in each micro-tract, per vascular zone. (E – I) The distributions of the 5 features for micro-tracts in each vascular zone overlap; the curves for each vascular zone are kernel density estimates. E and F quantify the main direction of the tract compared to the blood vessel. G and H quantify how much the tract wraps around the circumference of the blood vessel. I quantifies how much of the length is covered by the micro-tract. Violins show the distribution, the box the interquartile range and the white line the median. Vascular zones sharing a letter are not significantly different; those sharing no letter differ at adjusted p < 0.05 (Kruskal–Wallis p = 1.91e-06, 2.11e-04, 0.005, and 7.31e-12 for panels A – D, respectively; Dunn’s post-hoc with Bonferroni correction) Panel C lacks letters because all post-hoc comparisons had p > 0.05.

**Supplementary Figure 6.**
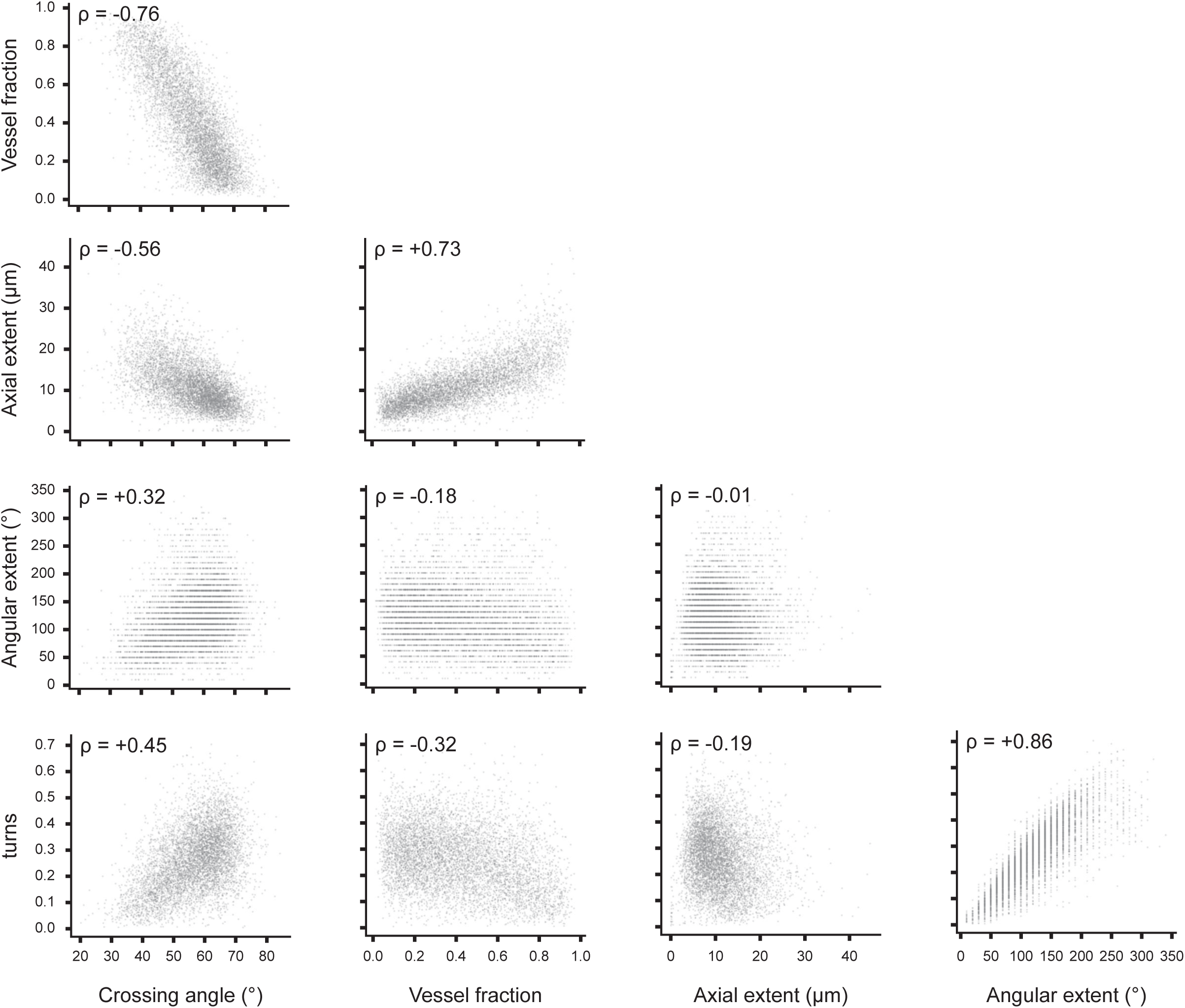
Correlations between micro-tract geometric features. Scatterplots of the 5 geometric micro-tract features against each other; ρ is Spearman’s correlation.

**Supplementary Figure 7.**
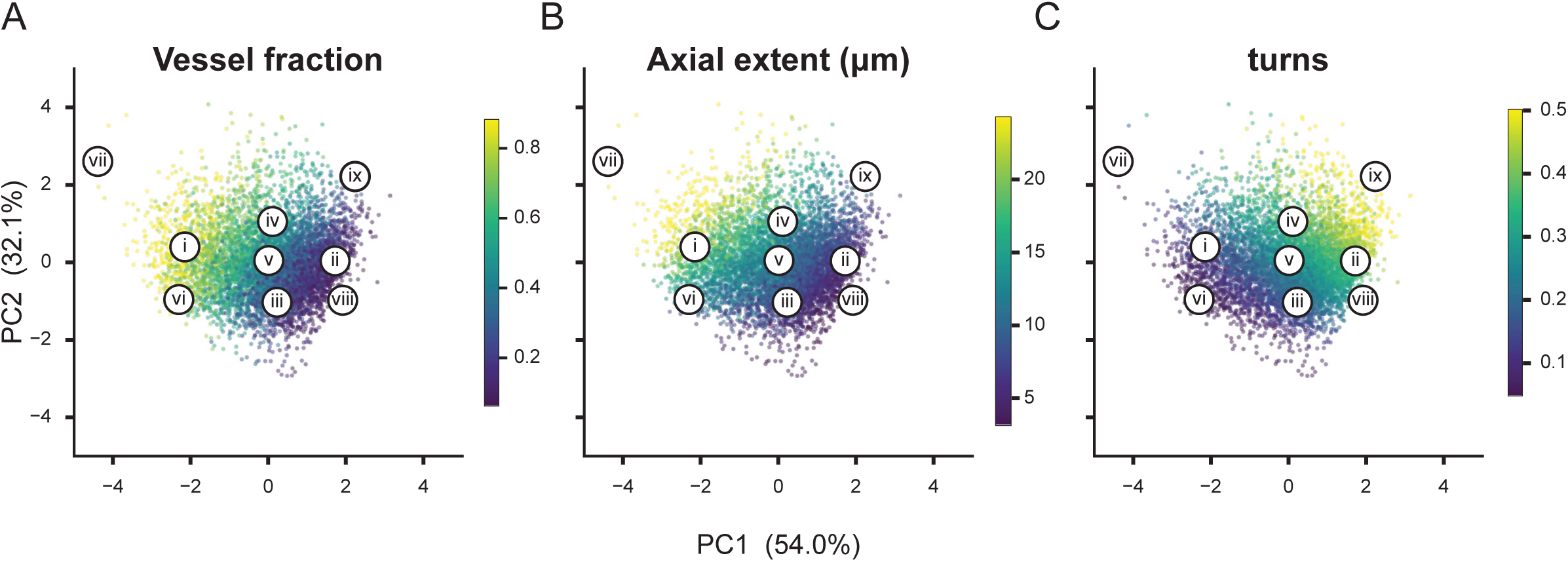
Micro-tract features in PCA space. As the loading on PC1 increases, micro-tracts (A) have less of their spatial variance along the vessel length, (B) occupy less of the length of the branch, and (C) rotate around the vessel more times. Roman numerals match the tracts shown in Figure 4F.

**Supplementary Figure 8.**
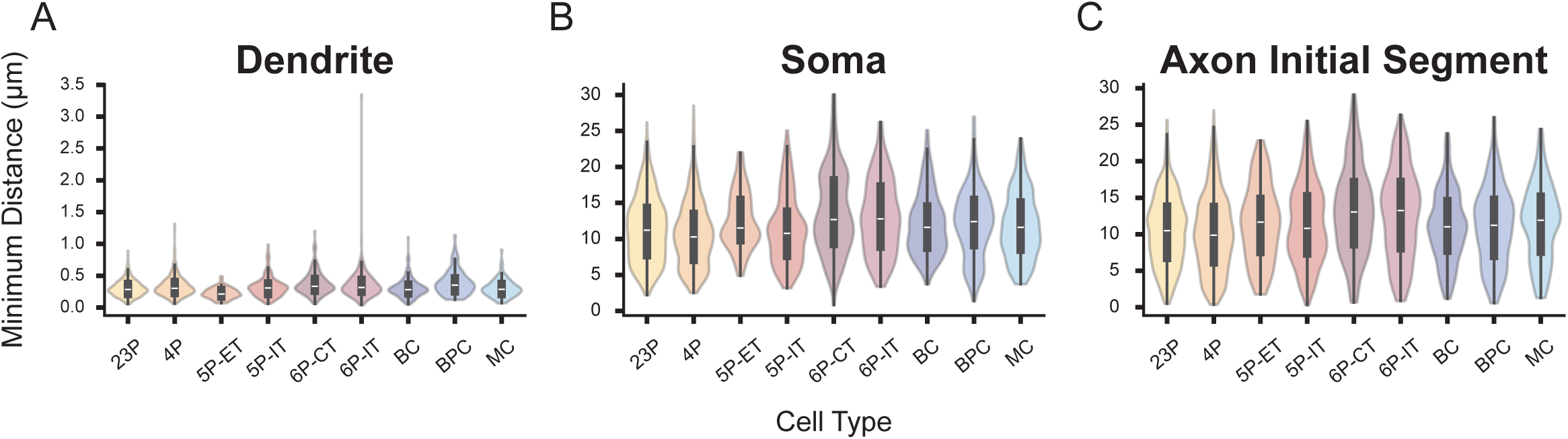
Spatial relationship between compartments of the proofread cells and the vascular network. (A – C) Minimum distance from any blood vessel to each cell’s (A) dendrites, (B) soma and (C) axon initial segment (AIS), by cell type. Violins show the distribution, the box the interquartile range and the white line the median. Note the different y-axis scale in A: dendrites came within a median of 0.2–0.4 μm of a vessel, while somata and AIS remained a median of 10–13 μm away. Differences among cell types within each panel were small, so compact letters are not shown; n per cell type in Supplementary Table 1.

**Supplementary Figure 9.**
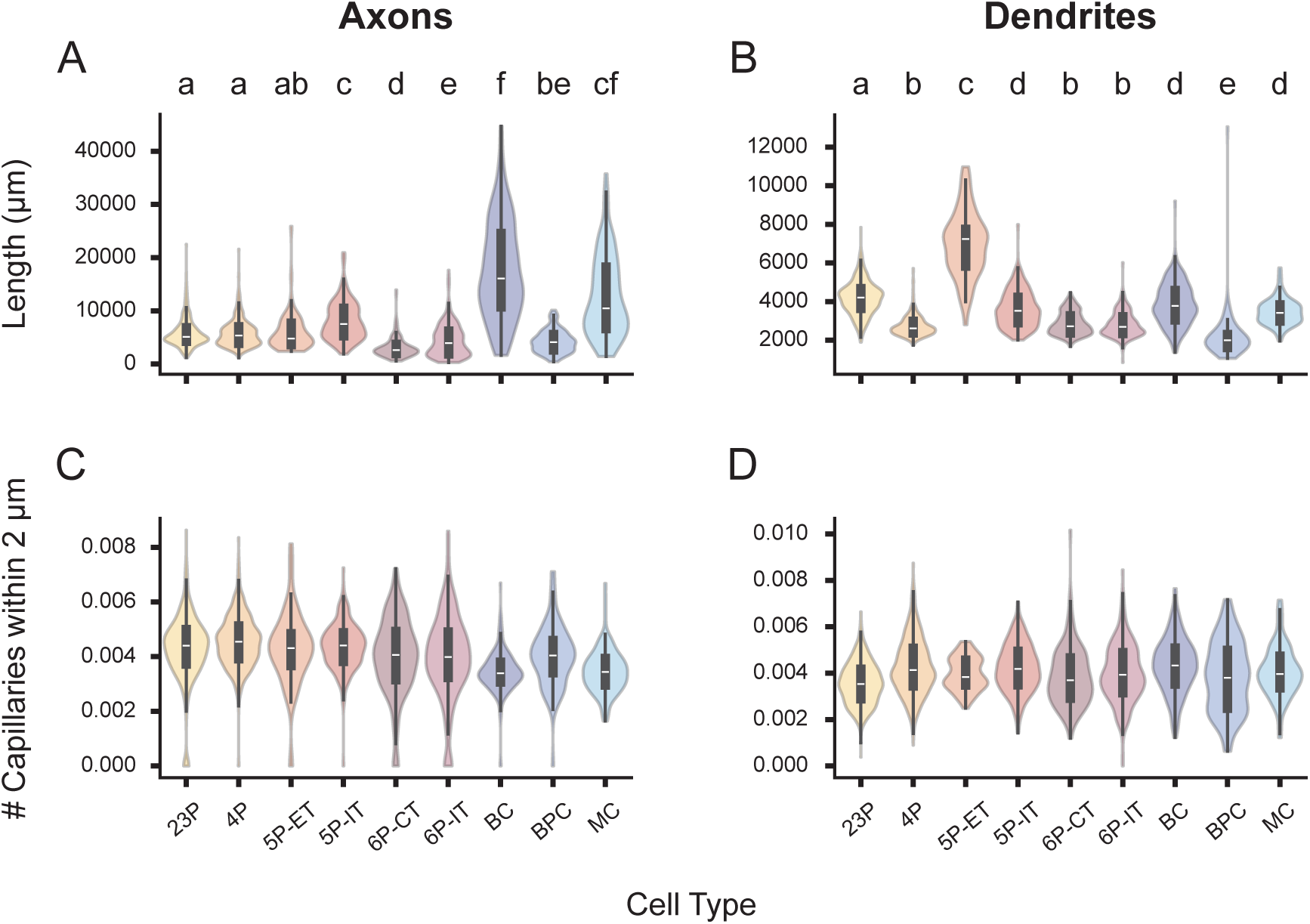
Cell types with longer axons and dendrites reach more capillaries. (A, B) Total (A) axon and (B) dendrite length per cell, by cell type. (C, D) Number of capillaries within 2 μm of each cell’s (C) axon and (D) dendrite, normalized by the total length of that process. Violins show the distribution, the box the interquartile range and the white line the median. In A and B, cell types sharing a letter are not significantly different; those sharing no letter differ at adjusted p < 0.05 (Kruskal–Wallis p = 4.3e-153 and 4.3e-224 respectively; Dunn’s post-hoc with Bonferroni correction). In C and D the medians span a narrow range around 0.004 capillaries per μm, so letters are not shown. n per cell type in Supplementary Table 1.

**Supplementary Table 1.** Median (IQR) number of distinct vessel branches of each class coming within 2 μm of a cell’s axon or dendrite, by cell type. n is the number of proofread neurons contributing to each row; axon and dendrite counts come from the same cells. A dash marks a class where the median and both quartiles were zero. The capillary column is shaded.

|  | Cell type | n | PA | ACT | term. PA | C | term. AV | CVT | AV | PCV | deep CVT | term. PCV | Pial |
| --- | --- | --- | --- | --- | --- | --- | --- | --- | --- | --- | --- | --- | --- |
| **Axon** | 23P | 536 | – | – | – | 22 (17–31) | 0 (0–1) | 1 (0–2) | 0 (0–1) | – | – | – | – |
|  | 4P | 573 | – | – | – | 23 (17–33) | – | 1 (0–2) | 0 (0–1) | – | – | – | – |
|  | 5P-ET | 65 | – | – | – | 20 (15–33) | – | 0 (0–1) | – | – | – | – | – |
|  | 5P-IT | 195 | 0 (0–1) | – | – | 33 (22–48) | 0 (0–1) | 1 (0–2) | 0 (0–1) | – | – | – | – |
|  | 6P-CT | 161 | – | – | – | 11 (7–16) | – | – | – | – | – | – | – |
|  | 6P-IT | 218 | – | – | – | 14 (6–26) | – | – | – | – | 0 (0–1) | – | – |
|  | BC | 210 | 0 (0–1) | – | – | 54 (35–84) | 1 (0–1) | 2 (0–4) | 1 (0–3) | – | – | – | – |
|  | BPC | 85 | – | – | – | 15 (10–21) | 0 (0–1) | 0 (0–1) | 0 (0–1) | – | – | – | – |
|  | MC | 112 | 0 (0–1) | 0 (0–1) | – | 34 (22–64) | 0 (0–1) | 2 (1–4) | 1 (0–3) | – | – | – | – |
| **Dendrite** | 23P | 536 | – | – | – | 14 (12–18) | – | 1 (0–2) | 0 (0–1) | – | – | – | – |
|  | 4P | 573 | – | – | – | 11 (9–14) | – | 0 (0–1) | 0 (0–1) | – | – | – | – |
|  | 5P-ET | 65 | – | – | – | 29 (21–32) | 0 (0–1) | 0 (0–1) | 0 (0–1) | – | – | – | – |
|  | 5P-IT | 195 | – | – | – | 15 (12–19) | – | 0 (0–1) | – | – | – | – | – |
|  | 6P-CT | 161 | – | – | – | 10 (8–14) | – | – | – | – | – | – | – |
|  | 6P-IT | 218 | – | – | – | 11 (8–14) | – | – | – | – | – | – | – |
|  | BC | 210 | – | – | – | 17 (12–21) | – | 0 (0–1) | 0 (0–1) | – | – | – | – |
|  | BPC | 85 | – | – | – | 8 (5–11) | – | 0 (0–1) | – | – | – | – | – |
|  | MC | 112 | – | – | – | 14 (11–17) | – | 0 (0–1) | – | – | – | – | – |

## Supplementary Notes

### Algorithm 1. Object clustering

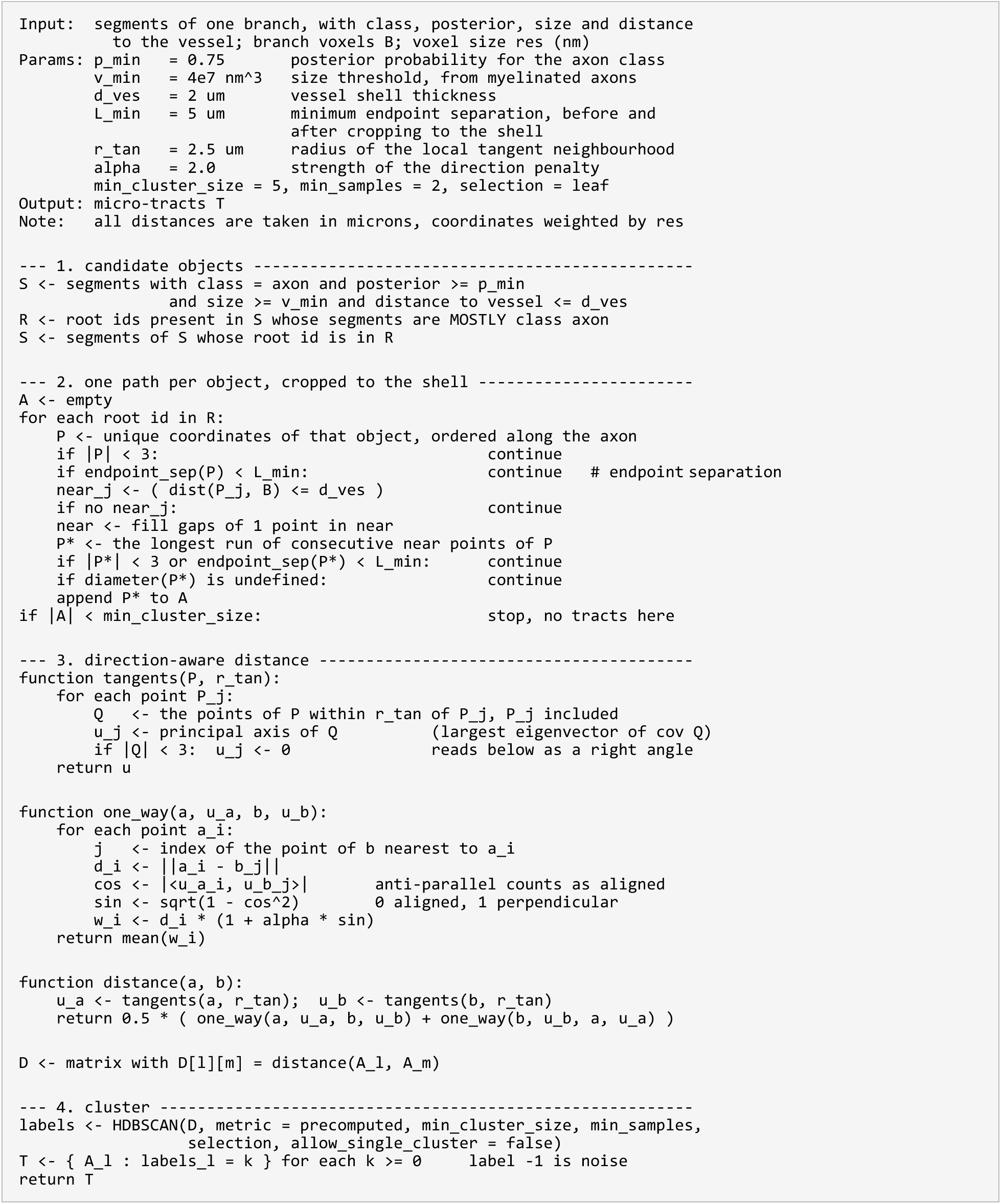

### Algorithm 2. Characterization of micro-tract geometry

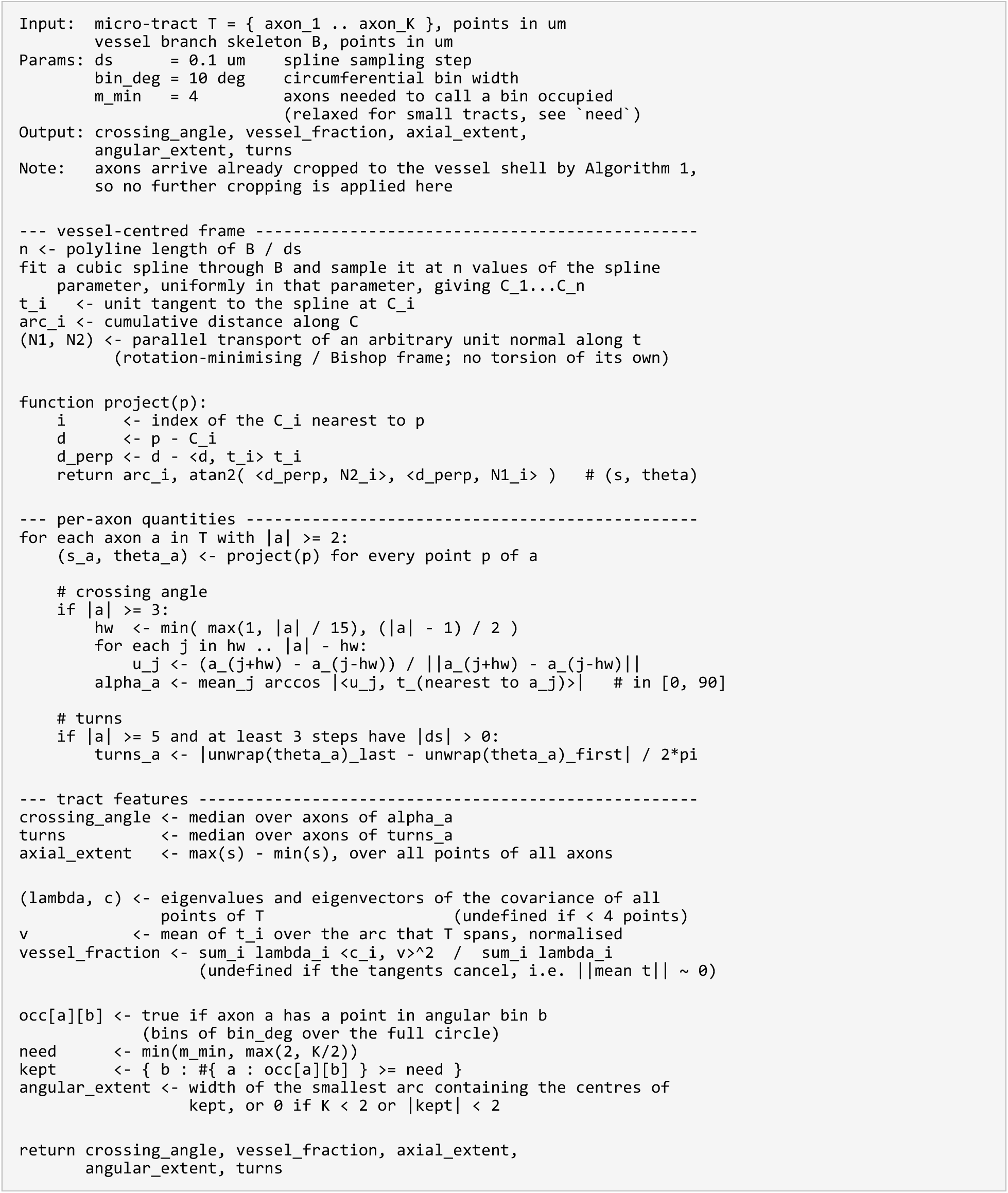

